# Modeling and Correction of Protein Conformational Disease in iPSC-derived Neurons through Personalized Base Editing

**DOI:** 10.1101/2024.01.17.576134

**Authors:** Colin T Konishi, Nancy Mulaiese, Tanvi Butola, Qinkun Zhang, Dana Kagan, Qiaoyan Yang, Mariel Pressler, Brooke G Dirvin, Orrin Devinsky, Jayeeta Basu, Chengzu Long

**Affiliations:** NYU Cardiovascular Research Center, NYU Grossman School of Medicine, New York, NY 100016; Leon H. Charney Division of Cardiology, NYU Grossman School of Medicine, New York, NY 100016; Department of Neuroscience and Physiology, NYU Grossman School of Medicine, New York, NY 100016; Department of Neurology, NYU Grossman School of Medicine, New York, NY 100016

**Keywords:** Protein conformational disease, protein aggregation, iPSC-derived neurons, CRISPR/Cas9, adenine base editing, engineered virus-like particles

## Abstract

Altered protein conformation can cause incurable neurodegenerative disorders. Mutations in *SERPINI1*, the gene encoding neuroserpin, can alter protein conformation resulting in cytotoxic aggregation leading to neuronal death. Familial encephalopathy with neuroserpin inclusion bodies (FENIB) is a rare autosomal dominant progressive myoclonic epilepsy that progresses to dementia and premature death. We developed HEK293T and induced pluripotent stem cell (iPSC) models of FENIB, harboring a patient-specific pathogenic *SERPINI1* variant or stably overexpressing mutant neuroserpin fused to GFP (MUT NS-GFP). Here, we utilized a personalized adenine base editor (ABE)-mediated approach to correct the pathogenic variant efficiently and precisely to restore neuronal dendritic morphology. ABE-treated MUT NS-GFP cells demonstrated reduced inclusion size and number. Using an inducible MUT NS-GFP neuron system, we identified early prevention of toxic protein expression allowed aggregate clearance, while late prevention halted further aggregation. To address several challenges for clinical applications of gene correction, we developed a neuron-specific engineered virus-like particle to optimize neuronal ABE delivery, resulting in higher correction efficiency. Our findings provide a targeted strategy which may treat FENIB and potentially other neurodegenerative diseases due to altered protein conformation such as Alzheimer’s and Huntington’s diseases.

## Introduction

Altered protein conformation can cause progressive and incurable neurodegenerative disorders such as Alzheimer’s and Huntington’s diseases.^1,2^ Neuroserpin (NS) is a neuron-specific serine protease inhibitor involved in axon growth and synapse formation.^3,4^ Some mutations in the *SERPINI1* gene alter neuroserpin protein stability, leading to protein aggregation within the endoplasmic reticulum (ER), causing ER and oxidative stress, neuronal dysfunction, and cell death.^5–7^ Familial encephalopathy with neuroserpin inclusion bodies (FENIB) is a rare autosomal dominant progressive myoclonic epilepsy, causing seizures, dementia, motor impairments, and premature death.^8–10^ The variant, *SERPINI1* c.1175 G>A (p.G392E) causes one of the most severe forms of FENIB, characterized by absence seizures, myoclonic and tonic-clonic seizures, and progressive dementia beginning in late childhood to early adolescence.^11,12^ There are no disease-modifying therapies.^13,14^ Many clinical and preclinical strategies for the treatment of conformational diseases, including Alzheimer’s and Huntington’s diseases, aim to reduce or eliminate expression of precursor forms of aggregation-prone proteins.^15,16^ These strategies employ RNAi, antisense oligonucleotides, small molecular modulators, or genome editors such as zinc-finger nucleases, transcription activator-like effector nucleases (TALEN), and clustered regularly interspace palindromic repeat (CRISPR)/CRISPR-associated sequence (Cas) nucleases to target pathogenic forms of RNA and DNA.^17–20^ All approaches face their own challenges.

CRISPR/Cas genome editors are sequence-specific RNA-guided nucleases that generate double-strand breaks (DSB) in target DNA.^21–23^ DSB repair through non-homologous end joining facilitates gene disruption due to the formation of insertions and deletions, while homology directed repair (HDR) produces sequence-specific modifications by introducing an exogenous repair template.^24,25^ Cas9 can edit stable cell lines and induced pluripotent stem cells (iPSCs) *in vitro* and neurons, cardiomyocytes, hepatocytes, and other cell types *in vivo*.^26–28^ Advances in iPSC technology and CRISPR/Cas genome editing have enabled the generation of isogenic cells harboring patient-specific disease-causing variants. These cells can be differentiated into various cell fates to model disease.^29–31^ However, HDR has been demonstrated to be primarily active during S and G2 phases of the cell cycle which limits its therapeutic applications in post-mitotic cells.^32–34^ Adenine base editing (ABE), a CRISPR/Cas9 modality, uses an evolved transfer RNA deaminase (tadA) fused to Cas9 nickase (nCas9) to act on single-stranded DNA, converting adenine (A) to guanine (G) without a DSB. ABEs can generate precise A>G transition mutations and have the potential to correct the pathogenic *SERPINI1* c.1175 G>A mutation.^35^

Therapeutic delivery of gene editors remains a common challenge for all gene therapies.^36^ Adenovirus, adeno-associated virus (AAV), and lentivirus are highly efficient viral vectors but have limited scope due to immunogenicity and genomic packaging capacity inherent to using viruses.^37,38^ Engineered virus-like particles (eVLPs) are an extracellular vesicle formed of viral proteins that can deliver cargo fused to retroviral Gag proteins but do not contain viral genomes. eVLPs target tissues with far fewer immunogenic effects and are expressed for a shorter duration than AAVs and lentivirus, thereby reducing off-target edits.^38–40^ Further, eVLPs can target cell-types by modulating surface glycoproteins.^39^ However, the efficiency of eVLP- delivered ABEs to correct mutations in cortical neurons remains unknown, as neurons use nonclassical nuclear import, with lower expression of KPNA2 and KPNA7, which are recognized by proteins with classical nuclear localization signals.

In this proof-of-concept study, we generated a stable overexpression model of FENIB in human embryonic kidney 293T (HEK293T) cells, and for the first time, endogenous knock-in and overexpression models of FENIB in iPSC-derived neurons. We used ABEs to target and correct the pathogenic variant. Correction of the variant restored dendritic morphology in KI neurons, and clearance of MUT NS-GFP aggregates in HEK293T cells. Using our inducible system, early treatment of MUT NS-GFP cells led to aggregate clearance, while later treatment prevented aggregate progression. We also tested eVLPs with neuron-specific nuclear localization signals (NLS) and identified improved neuronal editing. eVLP-mediated ABE editing appears to be a safe and efficient treatment for FENIB and potentially other diseases of protein conformation.

## Results

### CRISPR/Cas9 mediated HDR KI of patient-specific *SERPINI1* variant

The autosomal dominant *SERPINI1* c.1175 G>A (p. G392E) variant causes neuroserpin to form cytotoxic aggregates (**Fig. 1A, B**).^9^ To investigate ABE correction as a means to treat this variant, we first generated a *SERPINI1* c.1175 G>A cell line using CRISPR/Cas9 mediated HDR to knock-in the variant in HEK293T cells. We tested SpCas9 and SpCas9-NG editors to target *SERPINI1* exon 9 (**Fig. 1B**).^41^ Single guide RNAs (gRNA) were cloned into plasmids encoding SpCas9-2A-GFP (g1) or SpCas9-NG-2A-GFP (g2). g1 and g2 cleaved *SERPINI1* proximal to c.1175, and a single stranded oligodeoxynucleotide donor (ssODN) was used for templated HDR-mediated knock-in. g1 used a 5’-GGG-3’ PAM located at *SERPINI1* c.1173-c.1175. KI of *SERPINI1* c.1175 G>A alters the PAM sequence to 5’-GG<u>A</u>-3’, preventing recutting by SpCas9, to increase KI efficiency by preventing recurrent cleavage. g2 used a 5’-TG-3’ PAM to generate a DSB immediately downstream of the KI sequence to optimize HDR-mediated KI efficiency.^42^ First, HEK293T cells were transfected using lipofectamine 2000 with g1 or g2 plasmids without ssODN (**Fig. S1A, B**). 72-hours post-transfection, cells were sorted by GFP expression, and editing efficiency was measured by PCR-based T7 endonuclease I (T7E1) assay, Sanger sequencing, and Synthego Inference of CRISPR Edits (Synthego ICE) analysis tool.^43,44^ Both g1 and g2 demonstrated high editing activity on target sites. Next, we nucleofected HEK293T cells with g1 or g2 plasmids and *SERPINI1* c.1175 G>A ssODN for templated KI using the Lonza nucleofection SF system (**Fig. S1C**). 72-hours post-nucleofection, we sorted GFP-positive cells into bulk populations or single cells to assess editing efficiencies and selected clones harboring the patient-specific *SERPINI1* c.1175 G>A variant. ICE identified modest HDR efficiencies by g1 (16.5%) and g2 (11.5%). g2 produced far fewer nonspecific indels (50%) compared to g1 (80%). The similar KI efficiencies but improved nonspecific indel profile of g2 prompted us to focus on g2 for HEK293T modeling.

**Figure 1.**
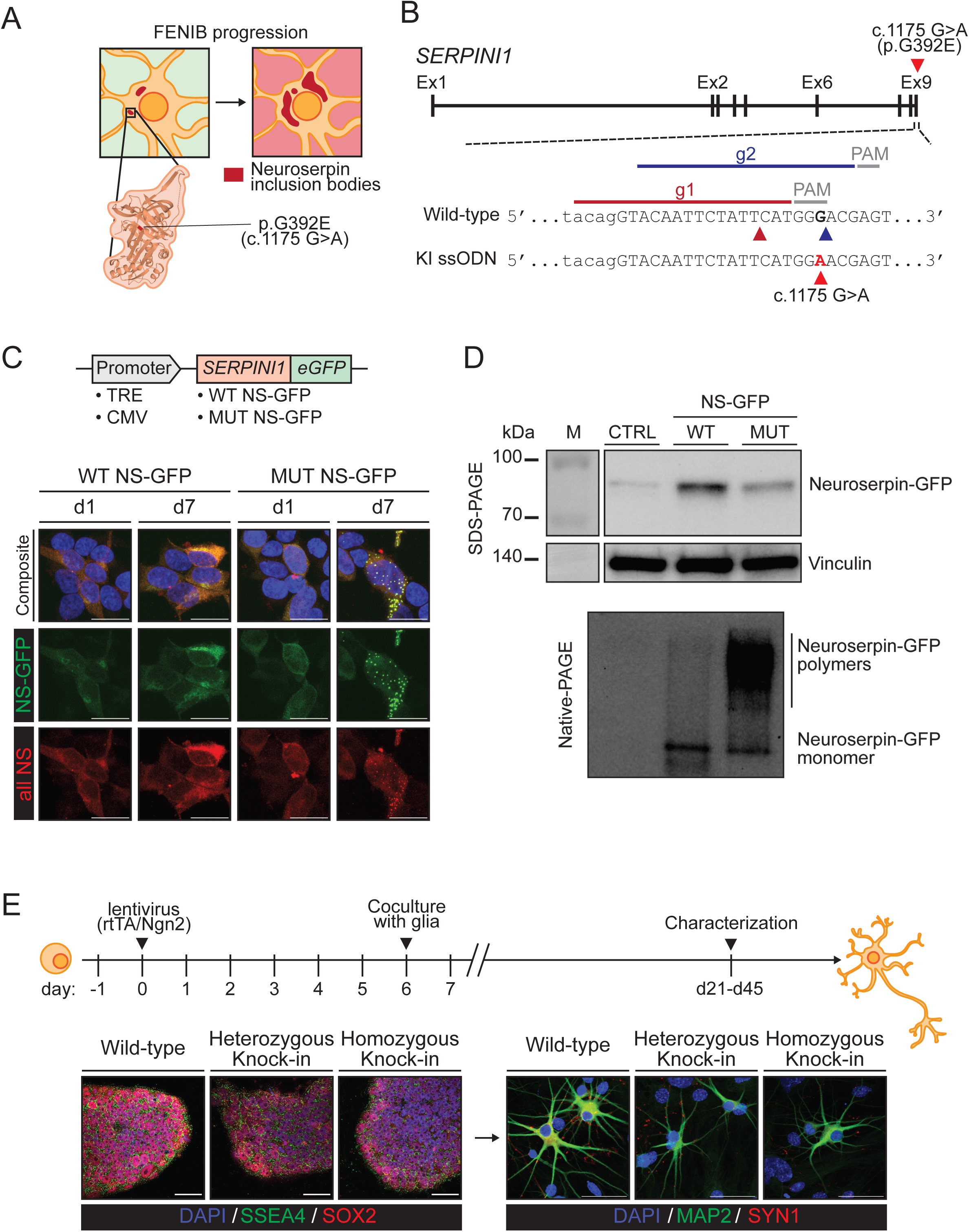
Generation of FENIB models. **(A)** FENIB disease progression. Mutant neuroserpin protein aggregates into neuroserpin inclusion bodies. **(B)** *SERPINI1* gene locus. Top: patient-specific *SERPINI1* c.1175 G>A (p.G392E) mutation located in exon 9 (red arrowhead). Middle: gRNAs for targeting wild-type *SERPINI1* (g1 in dark red and g2 in blue). PAM (grey) and DNA cleavage sites indicated with matching arrowheads. Bottom: knock-in (KI) single-stranded oligodeoxynucleotide (ssODN) sequence with patient-specific c.1175 mutation in red (red arrowhead). **(C)** Wild-type (WT) or mutant (MUT) neuroserpin fused to GFP (NS-GFP) lentivirus under the tetracycline response element (TRE) or cytomegalovirus (CMV) promoter used to generate stably expressed NS-GFP cells. Representative confocal microscopy images of HEK293T cells expressing WT NS-GFP or MUT NS-GFP under the TRE promoter. Samples imaged at day 1 (d1) and day 7 (d7) post-doxycycline induction and blasticidin selection. HEK293T cells express fluorescent NS-GFP (green) and are labeled with antibody specific for neuroserpin (all NS) (red). Scale bar, 25 µm. **(D)** Immunoblot of cell lysate from HEK293T cells expressing WT NS-GFP, MUT NS-GFP, or control. Samples were resolved using anti-neuroserpin antibody in SDS-PAGE (top), anti-vinculin antibody in SDS-PAGE as control (middle), or anti-neuroserpin antibody in native-PAGE (bottom) to detect polymers. **(E)** iPSCs transduced with lentivirus expressing rtTA and Ngn2. Ngn2 expression induced for 7 days using doxycycline with puromycin selection for 5 days. Cells replated onto mouse glia and cultured in neurobasal media containing neurotrophic factors. Representative confocal microscopy images of iPSCs and d21 neurons expressing stage-specific proteins. iPSCs are labeled with antibodies specific for stage-specific embryonic antigen 4 (SSEA4) (green) and SRY-Box transcription factor 2 (SOX2) (red). Neurons are labeled with antibodies for neuron specific microtubule associated protein 2 (MAP2) (green) and synapse protein synapsin I (SYN1) (red). Scale bar, 50 µm.

Next, we nucleofected iPSCs with either g1 or g2 plasmid and *SERPINI1* c.1175 G>A ssODN for templated KI. Both gRNAs efficiently generated cells harboring the variant sequence (g1: 24%, g2: 21%), consistent with our HEK293T cell editing (**Fig. S1D**). All identified FENIB patients are heterozygous for *SERPINI1* variants. ^45^ However, we sought to assess the impact of gene dosage and selected both heterozygous *SERPINI1* c.1175 G>A/+ (heterozygous KI) and homozygous *SERPINI1* c.1175 G>A/G>A (homozygous KI) clones. Patient-specific iPSCs and isogenic control wildtype clones were clonally expanded for future experiments.

### Stable overexpression of GFP fused neuroserpin in human cells

Previous studies have used immortalized cell lines, and mouse primary neural progenitors and neurons overexpressing neuroserpin protein to model FENIB.^5,14,46,47^ To more accurately recapitulate patient disease, we implemented overexpression of mutant neuroserpin in human HEK293T cells and iPSCs in addition to our KI cells. We designed a lentivirus vector expressing wild-type (WT) or mutant (MUT) neuroserpin protein fused to enhanced green fluorescent protein (NS-GFP) under constitutive cytomegalovirus (CMV) or doxycycline-inducible tetracycline response element (TRE) promoters (**Fig. 1C**). We transduced HEK293T cells with TRE-NS-GFP and reverse tetracycline-controlled transactivator (rtTA) lentiviruses. One day post-transduction, we induced NS-GFP expression by adding doxycycline to the culturing media. Using a combination of imaging native GFP fluorescence and immunocytochemical labeling, we assessed NS-GFP expression over the span of one week. One day after NS-GFP induction (d1), cells expressed NS-GFP with no observable differences between WT and MUT NS-GFP expressing cells. Both types of cells had diffuse NS-GFP fluorescence which colocalized with immunofluorescent neuroserpin staining. By day 7 (d7), MUT NS-GFP cells harbored aggregates in the form of concentrated puncta of GFP expression (**Fig. 1C**). Fluorescent GFP puncta colocalized with the anti-neuroserpin immunofluorescent staining. WT NS-GFP fluorescence remained cytoplasmic and diffuse throughout the duration of our experiments, with no detectable puncta at any timepoint.

Protein aggregation in MUT NS-GFP cells was confirmed using denaturing sodium dodecyl-sulfate polyacrylamide gel electrophoresis (SDS-PAGE) and nondenaturing native-PAGE immunoblotting (**Fig. 1D and Fig. S1E**). Immunoblot of SDS-PAGE using anti-neuroserpin antibody revealed a band of ∼85kDa in the WT and MUT NS-GFP lanes; the combined molecular weights of neuroserpin (55kDa) and GFP (30kDa) fusion protein (**Fig. 1D**). Native-PAGE immunoblots of WT and MUT NS-GFP confirmed observations made during fluorescent microscopy imaging. Native-PAGE immunoblots of WT NS-GFP revealed a single dominant band indicative of NS-GFP monomers. MUT NS-GFP samples had a NS-GFP monomer band in addition to banding and smearing at higher molecular weights, reflecting mutant NS-GFP aggregate polymers. Next, we validated our results by resolving SDS-PAGE and native-PAGE using anti-GFP antibodies (**Fig. S1E**). Immunoblot of SDS-PAGE using anti-GFP antibodies revealed a ∼85kDa band in both WT and MUT NS-GFP lanes. Validating results obtained with the anti-neuroserpin antibody, native-PAGE immunoblot of WT NS-GFP and MUT NS-GFP revealed a shared NS-GFP monomer band while MUT NS-GFP had additional high molecular weight banding and smearing.

Neuroserpin aggregation in the endoplasmic reticulum (ER) causes ER stress mediated through the unfolded protein response (UPR) and ER overload response (EROR).^7,48,49^ We collected RNA from WT and MUT NS-GFP HEK293T cells, cells stably expressing GFP, and control cells without any form of overexpression. We assessed ER stress through quantitative PCR (qPCR) of UPR response genes *BiP* and spliced *XBP1* (**Fig. S1F**). When normalized to control

HEK293T cells, cells overexpressing GFP show no difference in fold expression of either gene. We find that both WT and MUT NS-GFP cells revealed a significant increase in both *BiP* (WT NS-GFP: 4.1, MUT NS-GFP: 7.8, p<0.0001) and *sXBP1* (WT NS-GFP: 2.8, MUT NS-GFP: 3.9, p<0.0001), with the greatest increase in MUT NS-GFP cells when normalized to *18s* transcript expression. Previous studies of cell metabolic activity have found that overexpression of WT NS reduces cellular viability^14^. In all, our immunofluorescent microscopy, native-PAGE immunoblot, and transcript analyses demonstrated stable overexpression of NS-GFP results in FENIB neuroserpin aggregation in human cells.

### Generation of patient-specific iPSC-derived FENIB neurons through *Neurogenin2* induction

To evaluate our iPSC *in vitro* FENIB phenotype model, we differentiated iPSCs into excitatory cortical neurons by overexpressing *Neurogenin2* (Ngn2) (**Fig. 1E and Fig. S2A**).^31^ We assessed induction efficiency through immunocytochemical labeling of iPSCs prior to induction and cells 21 days (d21) post-induction. iPSCs showed standard iPSC morphology and pluripotency marker expression when labeled with stage-specific embryonic antigen 4 (SSEA4) and SRY-Box transcription factor 2 (SOX2). iPSC-derived neurons (iNs) displayed standard neuronal morphology and neuronal expression of neuron-specific microtubule associated protein-2 (MAP2), and synapse protein synapsin-1 (SYN1) (**Fig. 1E, Fig. S2B, and Fig. S2E**). At day 35 (d35), iNs also expressed neurofilament heavy (NFH) and neuronal nuclear protein (NeuN), characteristic of mature neurons (**Fig. S2C**).

Next, we sought to confirm the neural identity of iNs by comparing intrasample differential transcript expression in d35 iNs against previously published iN datasets (**Fig. S2D**).^31,50^ We collected RNA from three independent inductions per genotype and performed polyA selected bulk RNA-seq to target mRNAs. Transcriptional profiles of d35 neurons revealed an enrichment of genes confirming neuronal identity. In a comparison to the Zhang 2013 dataset, we identified pan-neuronal (*DCX*, *MAP2*, *TUBB3*), glutamate-related (*SLC17A6*), and post-synaptic (*DLG4*) genes within our top five hits. Similarly, comparing against the Lindhout 2020 dataset, we see further enrichment for axon (*GAP43*, *BASP1*), and neuron (*STMN1*, *ENO2*, *SYP*) genes. Further, we confirm that our iNs express *SERPINI1* (log_2_(TPM+1): wild-type: 7.14, heterozygous KI: 7.30, homozygous KI: 7.21, n = 3). Despite expression of *SERPINI1*, we were unable to detect NS aggregation in KI iNs throughout the duration of our studies. Our immunocytochemical and transcriptomic analyses together confirm the neural identity of Ngn2 induced neurons.

### iPSC-derived neuron models of FENIB demonstrate typical cortical neuron electrophysiology

We assessed the electrophysiological function of iPSC-derived neurons by analyzing evoked action potentials (AP) using whole-cell patch-clamp recordings in current clamp mode (**Fig. S3A-G**). While single APs were elicited in all neurons by day 14 (d14), with a burst of firing in a subset of tested cells, further maturation of iNs until day 42 (d42) resulted in consistent prototypical APs across all cells recorded. Hence, we assessed d42-44 iNs for intrinsic excitability. No significant differences were observed between wild-type, heterozygous KI, and homozygous KI neurons in AP rheobase (WT: 29.7 ± 2.1 pA, n = 54, HET.: 25.4 ± 1.6 pA, n = 46, HOM: 28.1 ± 2.0 pA, n = 47, p>0.05), full-width at half-maxima (WT: 2.0 ± 0.1 ms, n = 52, HET: 2.0 ± 0.1 ms, n = 46, HOM: 2.0 ± 0.1 ms, n = 47, p>0.05), amplitude (WT: 84.7 ± 1.2 mV, n = 52, HET: 82.5 ± 1.5 mV, n = 46, HOM: 84.6 ± 1.2 mV, n = 47, p>0.05), firing threshold (WT: -46.2 ± 1.1 mV, n = 52, HET: -48.5 ± 1.2 mV, n = 46, HOM:-47.9 ± 1.3 mV, n = 47 p>0.05), cell-capacitance (WT: 15.7 ± 0.7 pF, n = 53, HET: 15.5 ± 0.6 pF, n = 46, HOM: 16.5 ± 0.7 pF, n = 49,p>0.05), and resting membrane potential (WT: -49.9 ± 1.3 mV, n = 54, HET: -49.5 ± 1.3 mV, n = 45, HOM: -49.9 ± 1.6 mV, n = 50, p>0.05). Our evoked AP findings were consistent with *in vitro* cortical neurons, validating our human neuronal model of FENIB.^51–53^

### Characterizing neuroserpin inclusion bodies in NS-GFP overexpression iPSC-derived neuron FENIB model

We hypothesized that preventing mutant neuroserpin expression would facilitate clearance of pre-existing aggregated protein in MUT NS-GFP iNs (**Fig. 2A**).^54^ To test this, we transduced iPSCs with either doxycycline inducible WT or MUT NS-GFP lentivirus in addition to Ngn2 and rtTA lentivirus for inducing neuronal fate. Triply transduced cells were selected for following doxycycline induction using puromycin and blasticidin, with resistance conferred from Ngn2- T2A-PURO and NS-GFP-T2A-BSD lentivirus respectively.

**Figure 2.**
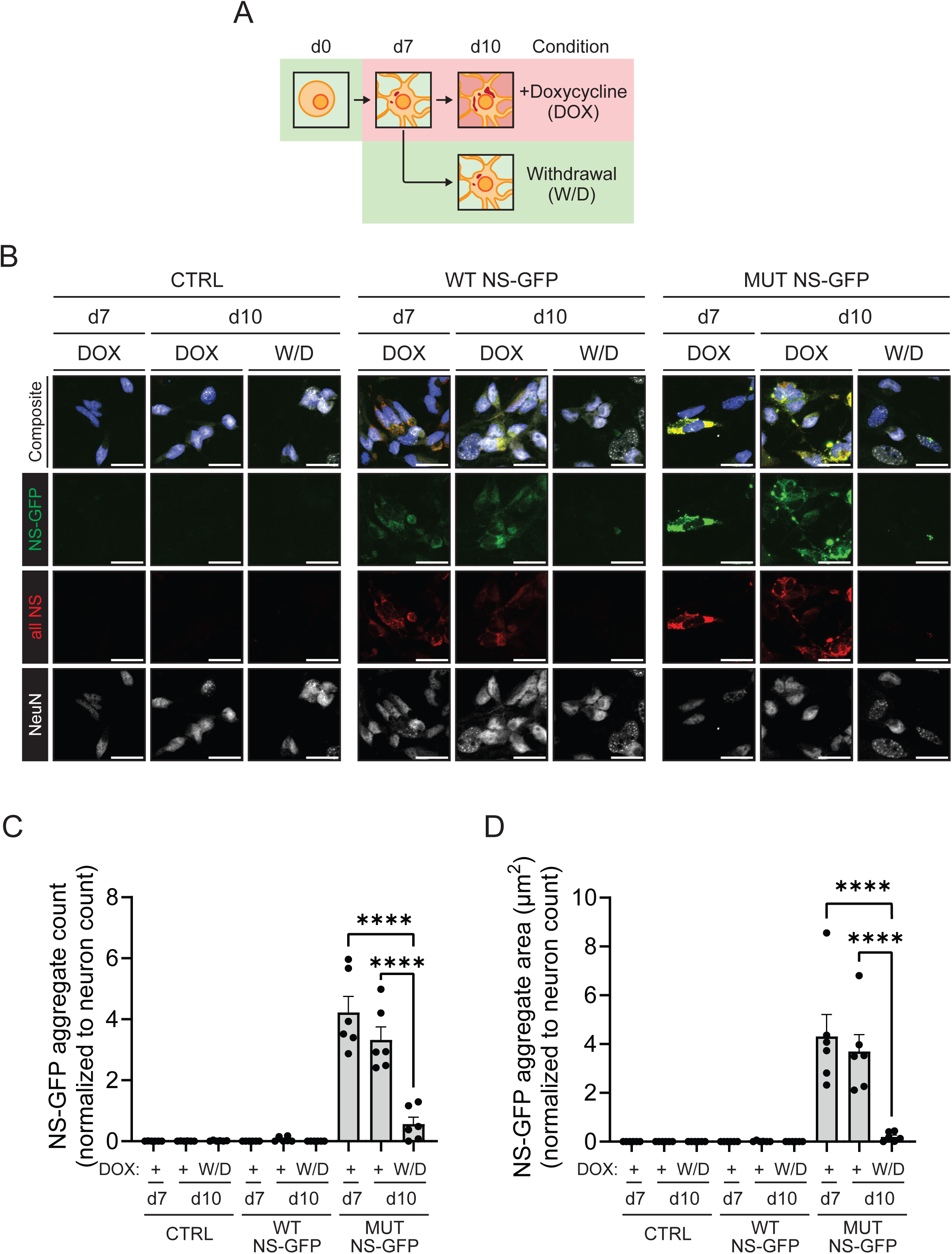
Early prevention of MUT NS-GFP expression is sufficient for aggregate clearance. **(A)** Administration and withdrawal of doxycycline for the aggregation and clearance of mutant NS-GFP. **(B-D)** Early prevention of MUT NS-GFP expression in inducible NS-GFP neurons. +Doxycycline (+DOX) cells received continuous doxycycline and expressed NS-GFP through day 10. Withdrawal (W/D) cells received doxycycline until day 7, at which point doxycycline was withdrawn, allowing aggregate clearance until day 10, at which point all cells were processed for imaging on day 10. Each sample is represented as a dot, indicative of an imaging field of a biological replicate. Each field has ∼100-200 cells. The graphs depict mean ± SEM of n = 6 biological replicates. **(B)** Representative confocal microscopy images of NS-GFP aggregation in day 7 and 10 FENIB neurons expressing WT NS-GFP, MUT NS-GFP, or control. Cells are labeled with antibodies specific for all neuroserpin forms (all NS) (red), NeuN (white), DAPI (blue), and express NS-GFP (green). Scale bar, 25 µm. **(C)** Quantification of NS-GFP aggregate count normalized to DAPI+/NeuN+ cells. (****p<0.0001) All comparisons left unmarked are statistically non-significant. **(D)** Quantification of NS-GFP aggregate area normalized to DAPI+/NeuN+ cells. (****p<0.001) All comparisons left unmarked are statistically non-significant.

NS-GFP expression was induced in iNs, with GFP signal observed one day after induction, similar to our HEK293T cell models. By d4, NS-GFP aggregates began forming in MUT NS-GFP cells. WT NS-GFP iNs behaved similarly to WT NS-GFP HEK293T cells, with NS-GFP signal detected as diffuse GFP. We sought to test whether early termination of mutant protein expression would facilitate the clearance of mutant protein aggregates. We induced expression of NS-GFP for 6 days. Beginning on d7, we separated cells into continued expression (+DOX) or discontinued expression through doxycycline withdrawal (W/D) until day 10 (**Fig. 2A**). We imaged d7 prior to separation, d10 +DOX, and d10 W/D cells using immunofluorescent confocal microscopy to identify aggregate formation and clearance in iNs (**Fig. 2B-D**). iNs were labeled with antibodies specific for NeuN and neuroserpin and expressed either WT NS-GFP, MUT NS-GFP, or a no overexpression control (**Fig. 2B**). Images were acquired as Z-stacks to capture fluorescent signal throughout the thickness of the iN culture. We then generated 2D reconstructions of the iNs using the “maximal projection” method in ImageJ to visualize the maximum signal of a given pixel in a 2D picture. d7 and d10 +DOX MUT NS-GFP cells harbored extensive NS-GFP aggregates in the form of GFP puncta which colocalized with neuroserpin immunofluorescent labeling. d7 and d10 +DOX WT NS-GFP cells had diffuse NS-GFP fluorescence which colocalized with neuroserpin immunofluorescent labeling. d10 W/D MUT NS-GFP cells revealed an almost complete clearance of NS-GFP aggregates. Similarly, d10 W/D WT NS-GFP cells had almost no detectable NS-GFP signal, which was verified by the loss of neuroserpin immunofluorescent labeling. Aggregated MUT NS-GFP signal produced distinct GFP puncta which are brighter than diffuse WT NS-GFP. Using ImageJ image analysis software, we set a fluorescence intensity threshold for GFP expression to establish an unbiased method to distinguish aggregated NS-GFP from diffuse NS-GFP signal. Next, we used the ImageJ particle analysis tool to quantify NS-GFP aggregate count and area **(Fig. S4A)**. These values were normalized to cells labeled with neuronal marker, NeuN, and 4’6-diamidino-2- phenylindole (DAPI) to obtain NS-GFP aggregate count per neuron and NS-GFP aggregate area per neuron. By day 7, MUT NS-GFP cells harbored extensive NS-GFP aggregates with no aggregates in either WT NS-GFP or no overexpression control (ctrl) counterpart cells (d7 MUT NS-GFP: 4.22 aggregates/neuron, d7 WT NS-GFP: 0 aggregates/neuron, ctrl: 0 aggregates/neuron, p<0.0001). Following doxycycline withdrawal, d10 W/D MUT NS-GFP cells had significantly reduced aggregate count compared to +DOX counterparts (d10 +DOX MUT NS-GFP: 3.32 aggregates/neuron, W/D MUT NS-GFP: 0.56 aggregates/neuron, p<0.0001). d10 W/D MUT NS-GFP values were statistically non-significant when compared to WT NS-GFP and no overexpression control aggregate counts (d10 +DOX ctrl: 0 aggregates/neuron, d10 W/D ctrl: 0 aggregates/neuron, d10 +DOX WT NS-GFP: 0.06 aggregates/neuron, W/D WT NS-GFP: 0 aggregates/neuron) (**Fig. 2C**). Aggregate clearance was also reflected in a reduction in aggregate area per neuron (d7 MUT NS-GFP: 4.31 µm^2^/neuron, d7 WT NS-GFP: 0 µm^2^/neuron, d7 ctrl: 0 µm^2^/neuron, p<0.0001). After doxycycline withdrawal, d10 W/D MUT NS-GFP cells had a statistically significant reduction in aggregate burden compared to continuously induced d10 +DOX MUT NS-GFP cells (d10 +DOX MUT NS-GFP: 3.70 µm^2^/neuron, d10 W/D MUT NS-GFP: 0.20 µm^2^/neuron, p<0.0001) (**Fig. 2D**). When compared to d10 WT NS-GFP and d10 no overexpression control counterpart cells, d10 W/D MUT NS-GFP aggregate area per neuron values were statistically non-significant (d10 +DOX ctrl: 0 µm^2^/neuron, d10 W/D ctrl: 0 µm^2^/neuron, d10 +DOX WT NS-GFP: 0.01 µm^2^/neuron, d10 W/D WT NS-GFP: 0 µm^2^/neuron). Based off these findings, early termination of mutant NS expression is sufficient for the clearance of aggregated protein.

Next, we assessed if prolonged aggregate accumulation could be cleared in more mature neurons (**Fig. 3A-D**). We induced aggregation until day 21 (d21), then split samples and used confocal microscopy to visualize and assess per neuron aggregation count and area at d21, day 28 (d28) (+DOX and W/D), and day 35 (d35) (+DOX and W/D) (**Fig. 3A**). Neurons were labeled with antibodies specific for NeuN and expressed MUT NS-GFP. (**Fig. 3B**). Using the ImageJ particle analysis method described above, we found that aggregate count remained similar from d21 to d28. W/D d28 iNs trended towards fewer aggregates, however, aggregate count between d21, +DOX d28, and W/D d28 iNs were not statistically significant (d21: 0.82 aggregates/neuron, +DOX d28: 0.71 aggregates/neuron, W/D d28: 0.62 aggregates/neuron, p>0.05) (**Fig. 3C**). When we terminated expression at d21 and assessed clearance in d35 neurons, W/D d35 iNs showed a significant decrease in aggregate count compared to +DOX counterparts (+DOX d35: 1.24 aggregates/neuron, W/D d35: 0.75 aggregates/neuron, p<0.01). W/D d35 aggregates per neuron was similar to d21 and d28 neurons (p>0.05). Next, we assessed aggregate area per neuron. W/D d35 iNs had significantly reduced aggregate area per neuron compared to +DOX iNs (+DOX d35: 3.2 µm^2^/neuron, W/D d35: 1.1 µm^2^/neuron, p<0.0001) (**Fig. 3D**). The difference in aggregate area per neuron identified between +DOX d35 iNs and W/D d35 iNs was not present when comparing W/D d35 iNs to d21 or either +DOX or W/D d28 iNs (d21: 1.427 µm^2^/neuron, +DOX d28: 1.16 µm^2^/neuron, W/D d28: 0.89 µm^2^/neuron, p>0.05). Thus, preventing MUT NS-GFP expression at day 21, and allowing cells to resolve toxic protein aggregation until day 35 was insufficient to clear aggregates. However, by stopping expression of the mutant protein, we were successful in preventing the formation of new aggregates. These doxycycline inducible NS-GFP studies demonstrate that early cessation of protein aggregation allowed for complete aggregate clearance. By contrast, late-stage cessation prevented further aggregate accumulation.

**Figure 3.**
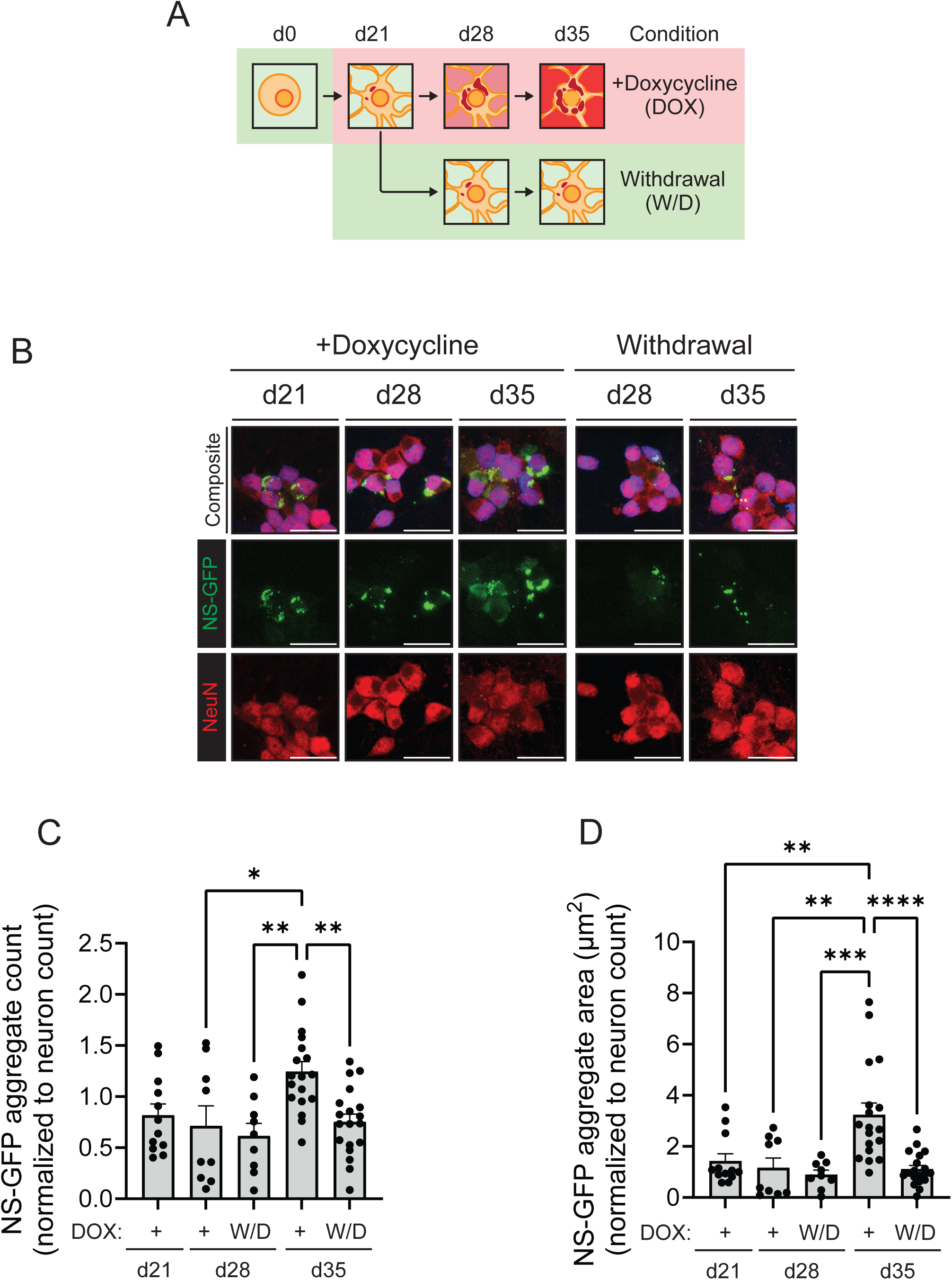
Late prevention of MUT NS-GFP expression prevents aggregation progression. **(A)** Administration and withdrawal of doxycycline for the aggregation and clearance of mutant NS-GFP. **(B-D)** Late prevention of MUT NS-GFP expression in inducible NS-GFP neurons. MUT NS-GFP aggregation in iPSC-derived neurons at indicated timepoints. Day 21, day 28, and day 35 neurons had continuous expression of MUT NS-GFP. Day 28 and day 35 withdrawal neurons had MUT NS-GFP expression withdrawn at day 21. Each sample is represented as a dot, indicative of an imaging field. Each field has ∼100-200 cells. The graphs depict mean ± SEM of n = 12-19 biological replicates (day 21 n = 12, day 28 dox and w/d n = 9, day 35 dox n = 18, day 35 w/d n = 19). **(B)** Representative confocal microscopy image of MUT NS-GFP neurons labeled with NeuN (red) and DAPI (blue), harboring NS-GFP aggregates (green). Scale bar, 25 µm. **(C)** Quantification of NS-GFP aggregate count. NS-GFP aggregate count is normalized to DAPI+/NeuN+ cells. (*p<0.05, **p<0.01). All comparisons left unmarked are statistically non-significant. **(D)** Quantification of NS-GFP aggregate area normalized to DAPI+/NeuN+ cells. (**p<0.001, ***p<0.001, ****p<0.0001). All comparisons left unmarked are statistically non- significant.

### iPSC-derived neurons overexpressing MUT NS-GFP show changes in membrane excitability

We next assessed neuronal excitability and physiological function using whole cell patch clamp in current clamp mode to evoke APs in NS-GFP iNs (**Fig. S5A-G**). Based on the maturation of neurons at day 42 (**Fig. S3**), NS-GFP iNs at d42 were assessed for intrinsic properties of excitability including rheobase, full-width at half-maxima, amplitude, firing threshold, cell capacitance, and resting membrane potential. +DOX MUT NS-GFP iNs had increased rheobase compared to the WT NS-GFP counterparts (+DOX MUT NS-GFP: 73.1 ± 14.3 pA, n = 8, +DOX WT MUT NS-GFP: 37.5 ± 5.9, n = 8, p<0.05) **(Fig. S5B)**. This suggests a decrease in membrane excitability consistent with other reports of protein aggregation based neurodegenerative disease models. ^55,56^ W/D d42 MUT NS-GFP (51.4 ± 5.1 pA, n = 7) neurons had similar rheobase as WT NS-GFP (+DOX WT NS-GFP: 37.5 ± 5.9 pA, n = 8, and W/D WT NS-GFP: 48.6 ± 7.0 pA, n = 7, p>0.05) and control (+DOX CTRL: 52.5 ± 4.5 pA, n = 8, and W/D CTRL: 63.3 ± 8.2, n = 9, p>0.05) neurons indicating that alteration to rheobase was rescued following termination of MUT NS-GFP expression. AP full-width at half-maxima (+DOX CTRL:1.6 ± 0.2 ms, n = 8, +DOX WT NS-GFP: 1.6 ± 0.1 ms, n = 8, +DOX MUT NS-GFP: 1.4 ± 0.1 ms, n = 8, W/D CTRL: 1.8 ± 0.2 ms, n = 9, W/D WT NS-GFP: 1.7 ± 0.2 ms, n = 7, W/D MUT NS-GFP: 1.6 ± 0.1 ms, n = 7, p>0.05), amplitude (+DOX CTRL: 87.0 ± 2.8 mV, n = 8, +DOX WT NS-GFP: 85.3 ± 3.3 mV, n = 8, +DOX MUT NS-GFP: 84.3 ± 2.0 mV, n = 8, W/D CTRL: 91.3 ±2.2 mV, n = 9, W/D WT NS-GFP: 85.6 ± 0.8 mV, n = 7, W/D MUT NS-GFP: 88.0 ± 3.6 mV, n = 7, p>0.05), firing threshold (+DOX CTRL: -49.0 ± 2.7 mV, n = 8, +DOX WT NS-GFP: -52.3 ± 1.6 mV, n = 8, +DOX MUT NS-GFP: -44.3.3 ± 2.9 mV, n = 8, W/D CTRL: -49.1 ± 2.7 mV, n = 9, W/D WT NS-GFP: -46.2 ± 2.8 mV, n = 7, W/D MUT NS-GFP: -48.8 ± 3.1, n = 7, p>0.05), cell capacitance (+DOX CTRL: 30.2 ± 3.9 pF, n = 8, +DOX WT NS-GFP: 24.4 ± 1.6 pF, n = 8, +DOX MUT NS-GFP: 26.1 ± 2.9 pF, n = 7, W/D CTRL: 27.2 ± 3.2 pF, n = 9, W/D WT NS-GFP: 24.0 ± 2.7 pF, n = 8, W/D MUT NS-GFP: 29.8 ± 4.2 pF, n = 6, p>0.05), and resting membrane potential (+DOX CTRL: -54.1 ± 2.1 mV, n = 11, +DOX WT NS-GFP: -54.2 ± 4.7 mV, n = 6, +DOX MUT NS-GFP: -56.6 ± 2.9, n = 7, W/D CTRL: -59.5 ± 2.5 mV, n = 8, W/D WT NS-GFP: -57.6 ± 5.0 mV, n = 8, W/D MUT NS-GFP: -61.9 ± 3.3 mV, n = 9, p>0.05) were all similar, with no statistically significant differences identified.

Based on our finding of dysfunctional membrane excitability due to an increase in rheobase in +DOX MUT NS-GFP iNs, we plotted an I-O curve to assess firing rate in +DOX and W/D iNs (**Fig. S5H and Fig. S5I**). When focused in on the minimum current required to increase the firing rate, we identified a minor trending but statistically non-significant difference between the initial increase in firing rate of d42 +DOX MUT NS-GFP iNs (40 pA) compared to WT NS-GFP (20 pA) and CTRL (30 pA) counterparts. Comparisons of d42 W/D iNs revealed no statistically significant differences. Taken together, our results demonstrate that MUT NS-GFP iNs have dysfunctional membrane excitability, which is rescued once MUT NS-GFP expression is terminated.

### ABE-mediated correction of *SERPINI1* c.1175 G>A variant

After identifying that termination of mutant neuroserpin expression through withdrawal of doxycycline in inducible MUT NS-GFP neurons led to aggregate clearance or prevented further aggregation, we sought to generate an ABE approach to target and correct the pathogenic variant, *SERPINI1* c.1175 G>A, back to the wild-type sequence. We identified a shortened 5’- TG-3’ PAM sequence downstream of the patient variant, recognized by ABEs with an SpCas9- NG C-terminal domain, which positioned the variant within the ABE editing window at the A5 position (**Fig. 4A**). Rationally designed and evolved ABE variants increase editing efficiency and reduce off-target effects.^57–59^ We tested four ABEs to identify an optimal ABE for targeting the *SERPINI1* c.1175 locus. We subcloned the SpCas9-NG C-terminal region into four next- generation ABEs resulting in NG-ABEmax, NG-SECURE-miniABEmax-V82G (NG-V82G), NG- SECURE-miniABEmax-K20AR21A (NG-K20AR21A), and NG-ABE8e.^58,60,61^ We found a candidate bystander mutation within the editing window at the A6 position. We hypothesized that this bystander mutation would be tolerated due to its location at the codon wobble position. Combination target and bystander editing encodes the wild-type residue (G, glycine), while bystander-only editing would not alter the pathogenic residue (E, glutamic acid).

**Figure 4.**
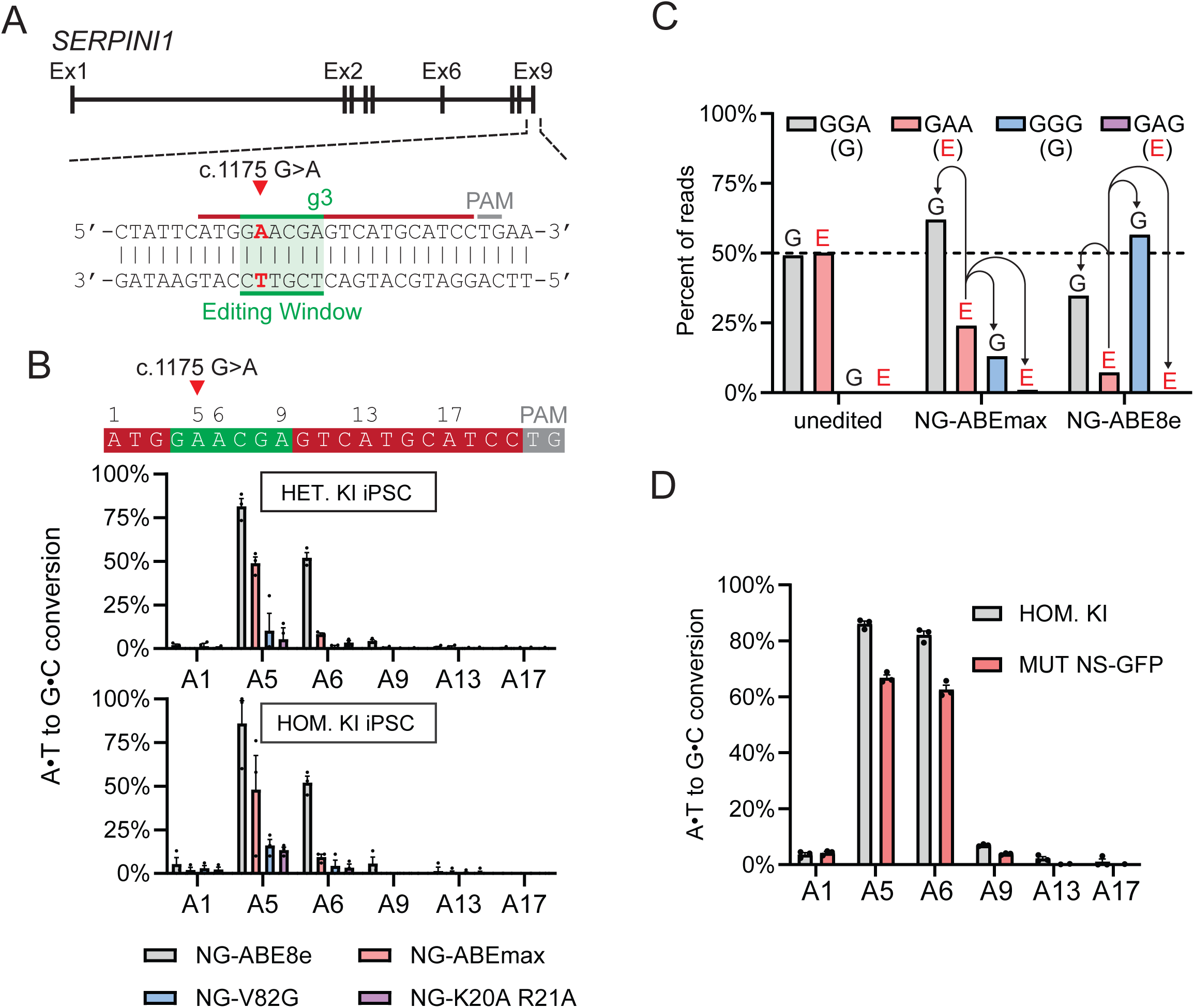
Correction of FENIB mutation in iPSC models using NG-ABE8e. **(A)** Patient-specific FENIB mutation *SERPINI1* c.1175 G>A targeted with adenine base editor (ABE). ABE editing window (green) using gRNA g3 (dark red) with NG PAM (grey). Patient-specific c.1175 G>A mutation is located at the A5 position of the gRNA sequence. **(B)** Quantification of ABE editing in heterozygous KI iPSCs (top) and homozygous KI iPSCs (bottom) using Sanger sequencing and EditR. The graphs depict mean ± SEM of n = 3 replicates. All adenine positions within the gRNA sequence are listed, with the c.1175 variant at position A5 (red arrowhead). **(C)** Quantification of deep sequencing reads of heterozygous KI iPSCs treated with NG-ABEmax or NG-ABE8e. Arrows represent editing outcomes from the variant GAA codon. Wild-type GGG codon (grey) encoding glycine (G; black), patient-specific variant GAA codon (red) encoding glutamic acid (E; red), wild-type synonymous GGG codon (blue) encoding glycine (G; black), and bystander only GAG codon (purple) encoding glutamic acid (E; red) are depicted above the graph. **(D)** Quantification of ABE editing using NG-ABE8e in homozygous KI and MUT NS-GFP model of FENIB. The graphs depict mean ± SEM of n = 3 replicates.

We cloned a mutant-specific sgRNA into the plasmid, pBSU6-sgRNA-GFP (g3), which was nucleofected along with a plasmid encoding ABE using the Lonza P3 nucleofection system in both heterozygous and homozygous KI iPSCs (**Fig. 4B**). We quantified ABE correction using EditR Sanger sequencing trace analysis and deep sequencing of the target locus compared to unedited control cells.^62^ EditR analysis of heterozygous KI iPSCs revealed the greatest correction efficiency with NG-ABE8e (82% A:T to G:C conversion efficiency) and NG-ABEmax (49%), and modest editing by NG-V82G (10%) and NG-K20AR21A (5%) at the A5 position. Analysis of homozygous KI iPSCs largely mirrored editing in heterozygous KI iPSCs (NG- ABE8e: 86%, NG-ABEmax: 48% NG-V82G: 16%, and NG-K20AR21A: 13% A:T to G:C conversion efficiency). Individuals harboring the *SERPINI1* c.1175 G>A variant are heterozygous as FENIB variants are often de novo mutations.^10–12^ We performed deep sequencing on heterozygous KI iPSCs treated with the two most efficient ABEs, NG-ABEmax and NG-ABE8e (**Fig. 4C**). Deep sequencing reads revealed codon specific editing outcomes with NG-ABEmax encoding 75% wild-type glycine (GGA or GGG) and NG-ABE8e encoding 92% glycine. Further analysis of glycine encoding reads revealed that NG-ABEmax activity resulted in more reversion to the wild-type codon (GGA: 62%, GGG: 13%) while NG-ABE8e had more codons harboring the permissive bystander mutation (GGA: 35%, GGG: 57%).

We treated MUT NS-GFP iPSCs with plasmids encoding g3 and NG-ABE8e to assess if NG- ABE8e editing could correct MUT NS-GFP iPSCs (**Fig. 4D**). EditR base calling analysis of Sanger sequencing traces revealed highly efficient editing in both homozygous KI (90%) and MUT NS-GFP (65%) iPSCs. Thus, NG-ABE8e with g3 can efficiently correct the variant within the endogenous genomic context in heterozygous and homozygous KI cells, and as the stably incorporated MUT NS-GFP model of FENIB.

### ABE correction of MUT NS-GFP in HEK293T cells allows aggregate clearance

After identifying a highly efficient method of base editing the pathogenic variant back to the wild-type *SERPINI1* sequence, we evaluated ABE treatment on NS-GFP aggregate clearance. Using lentivirus transduction, we stably incorporated either WT or MUT NS-GFP constitutively expressed under the CMV promoter into HEK293T cells. 72-hours after transduction, we sorted GFP positive cells into single cells to generate biological replicates. Cells were clonally expanded and expressed WT NS-GFP or MUT NS-GFP control for 3 weeks to induce extensive NS-GFP aggregation. MUT NS-GFP HEK293T clones were transfected with NG-ABE8e-T2A- tagRFP and g3 plasmids co-expressing tagRFP (ABE + g3), or NG-ABE8e-T2A-tagRFP only (ABE only) as control. 72-hours post-transfection, GFP+/RFP+ double positive cells were either bulk or single cell sorted to assess DNA correction and NS-GFP aggregate clearance. EditR analysis of bulk sorted cells revealed highly efficient correction of the variant back to the wild-type sequence in ABE + g3 MUT NS-GFP cells (71% A:T to G:C conversion efficiency) (**Fig. 5A**). We sought to visualize NS-GFP expression and aggregation using confocal microscopy in ABE + g3 MUT NS-GFP cells, and ABE only MUT NS-GFP, MUT NS-GFP pretreatment, and WT NS-GFP control cells (**Fig. 5B**). Native NS-GFP fluorescence was imaged alongside immunocytochemical labeling of neuroserpin using an antibody which recognizes all neuroserpin forms (1A10, all NS) and an antibody specific for aggregated neuroserpin polymers (purified polyclonal antibody, MUT poly). ^63^ We used Cellpose to count DAPI stained nuclei and determined total cells per imaging field to assess for culturing fitness differences. We did not identify any statistically significant changes (WT NS-GFP: 224 cells, pretreatment MUT NS-GFP: 235 cells, ABE only MUT NS-GFP: 233 cells, ABE + g3 MUT NS-GFP: 190 cells, p>0.05) (**Fig. 5C**). Immunoblotting whole cell lysate from ABE + g3 MUT NS-GFP, or ABE only MUT NS-GFP and WT NS-GFP controls revealed similar levels of NS-GFP protein expression when probed with anti-neuroserpin antibody (**Fig. S6A**). Therefore, any differences observed in aggregated and diffuse NS-GFP expression was caused by protein aggregation, as opposed to total protein expression.

**Figure 5.**
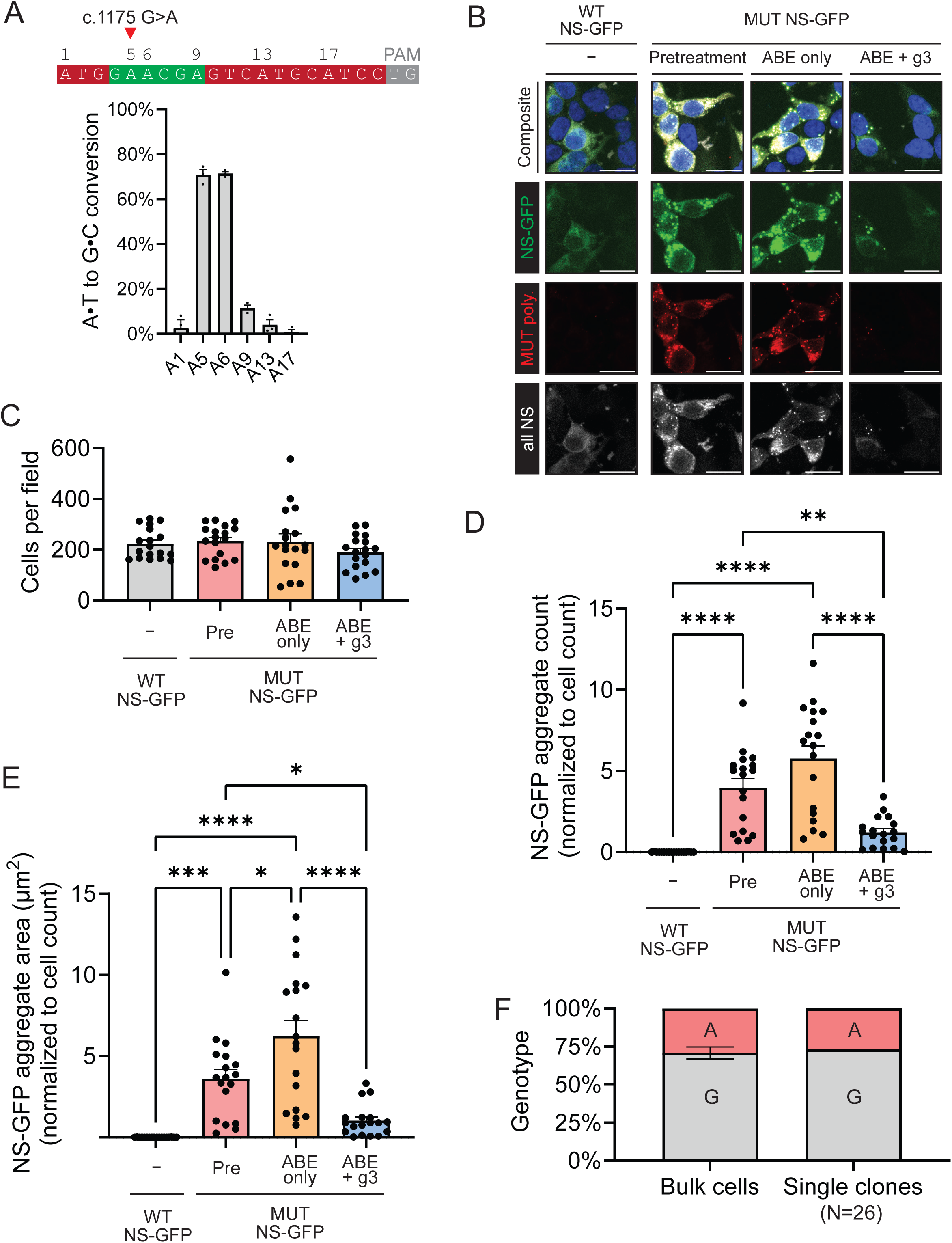
Correction of MUT NS-GFP in HEK293Ts allows for aggregate clearance. **(A)** Quantification of NG-ABE8e editing in MUT NS-GFP HEK293T cells using Sanger sequencing and EditR. gRNA sequence outlines targetable adenine bases, with the variant located at A5. The graph depicts mean ± SEM of n = 3 replicates. **(B)** Representative confocal microscopy images of ABE + g3 treated MUT NS-GFP, ABE only sham MUT NS-GFP control, pretreatment MUT NS-GFP control, or WT NS-GFP control HEK293T cells. Cells are labeled with antibodies specific for all neuroserpin forms (all NS) (grey), mutant neuroserpin polymers (MUT poly.) (red), DAPI (blue) and are expressing NS-GFP (green). Scale bar, 25 µm. **(C) Quantification of cells per field. All comparisons are statistically non-significant**. **(D-E)** Analysis of NS-GFP aggregate clearance following ABE + g3 treatment in confocal microscopy imaged coverslips**. (D)** Quantification of NS-GFP aggregate count normalized to cell count (****p<0.0001, **p<0.01). All comparisons left unmarked are statistically non-significant. **(E)** Quantification of NS-GFP aggregate area normalized to cell count (****p<0.0001, ***p<0.001, *p<0.05). All comparisons left unmarked are statistically non-significant. Each sample is represented as a dot, indicative of an imaging field. Each field has ∼200-400 cells. The graphs depict mean ± SEM of n = 3 biological replicates and n = 6 images per replicate. Pre = pretreatment. **(F)** Quantification of NG-ABE8e editing efficiency in MUT NS-GFP single clones (n = 26) following ABE + g3 treatment compared to bulk sorted cells using Sanger sequencing and EditR.

Next, we quantified the effects of ABE + g3 treatment of MUT NS-GFP cells on NS-GFP aggregation (**Fig. 5D and 5E**). ABE + g3 MUT NS-GFP cells had statistically significant reduced aggregates per cell values compared to pretreatment MUT NS-GFP and ABE only MUT NS-GFP cells (pretreatment MUT NS-GFP: 3.98 aggregates/cell, p<0.01, ABE only MUT NS-GFP: 5.77 aggregates/cell, p<0.0001, ABE + g3 MUT NS-GFP: 1.22 aggregates/cell) (**Fig. 5D**). ABE + g3 treatment was statistically non-significant when compared to WT NS-GFP control cells (WT NS-GFP: 0 aggregates/cell). ABE + g3 MUT NS-GFP cells also had a statistically significant reduction in NS-GFP aggregate area per cell compared to pretreatment MUT NS-GFP and ABE only MUT NS-GFP cells (pretreatment MUT NS-GFP: 3.61 µm^2^/cell, p<0.05, ABE only MUT NS-GFP: 6.23 µm^2^/cell,p<0.0001, ABE + g3 MUT NS-GFP: 1.02 µm^2^/cell) (**Fig. 5E**). ABE + g3 treatment was sufficient to restore aggregate area per cell back to WT NS-GFP (WT NS-GFP: 0 µm^2^/cell, p>0.05).

Next, we analyzed ABE + g3 treated single clones from two MUT NS-GFP HEK293T clones, Clone A and Clone B (**Fig. 5F and Fig. S6B**). Sanger sequencing of ABE + g3 treated single clones revealed that all clones (n = 26 total, Clone A = 6, Clone B = 20) were either corrected to the wild-type sequence (c.1175 G, Clone A = 3/6, Clone B = 16/20), or remained unedited (c.1175 G>A, Clone A=3/6, Clone B = 4/20), with no partially edited cells. Quantification of NG-ABE editing in single clones reflected editing efficiencies previously identified in bulk sorted cells (single clones: 72%, bulk sorted cells: 71%) (**Fig. 5F**). Following Sanger sequencing, we used epifluorescence microscopy to detect NS-GFP aggregates in ABE + g3 treated single clones (**Fig. S6B**). All corrected clones with the wild-type (c.1175 G) sequence had diffuse NS-GFP expression, while all unedited clones harbored NS-GFP aggregates. Based on our results, we conclude that ABEs can correct the FENIB mutation and allow aggregate clearance following extensive NS-GFP aggregation. Further, our clonal assay demonstrates that ABE correction is sufficient for total aggregate clearance in single cells.

### Impaired dendritic arborization in heterozygous and homozygous KI iPSC-derived neurons is partially restored with ABE treatment

Neuroserpin is expressed during late stage neuronal maturation with roles in dendritic morphology, synaptic maturation, and neuronal differentiation.^64–66^ Cultured rat hippocampal neurons overexpressing neuroserpin have increased dendritic arborization and altered spine shapes.^65^ *In vivo* analysis of neuroserpin-deficient mice revealed a decrease in spine- synapses.^67^ We evaluated whether dendritic morphology was affected in heterozygous and homozygous KI neurons, and whether ABE corrected neurons showed restored morphology **(Fig. S7A)**. We transfected d35 WT, heterozygous KI, homozygous KI, and ABE corrected neurons with GFP expressing plasmids for 30 minutes to sparsely label single cells. 24-hours post-transfection, we fixed coverslips for staining with anti-GFP antibodies to increase signal in imaging for Sholl analysis. Using ImageJ, we set a fluorescence intensity threshold against GFP expression to image the morphology of single neurons. Then using Simple Neurite Tracer (SNT), we assessed the number of dendritic intersections at 10 µm radial distances from the neuron soma **(Fig. 6A).**^68,69^

**Figure 6.**
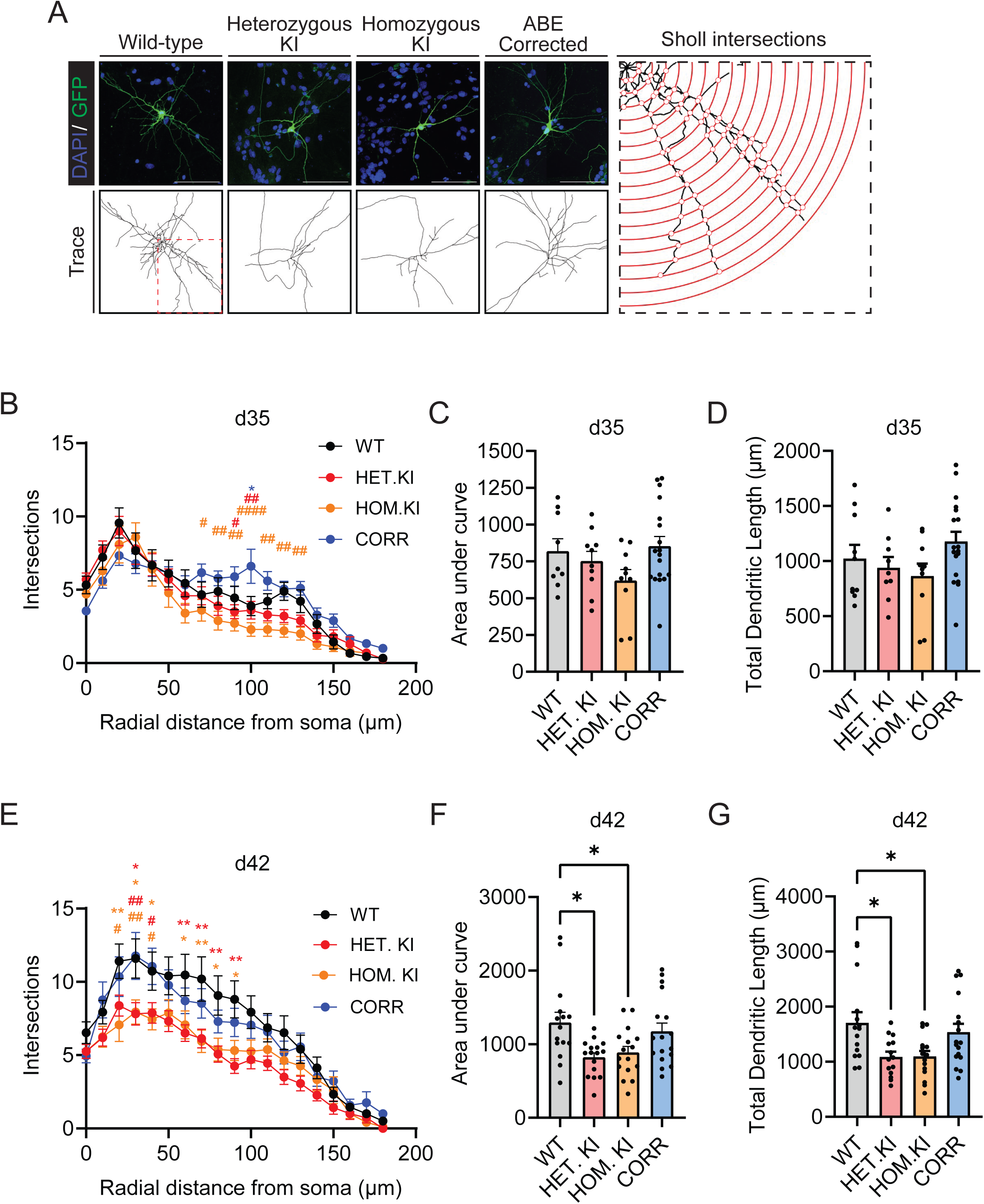
Dendritic outgrowth is impaired in FENIB KI neurons and is partially restored following ABE treatment. **(A)** Representative confocal microscopy images of d42 neurons labeled with GFP (green) and DAPI (blue), reconstructed neuron trace used for Sholl analysis, and representative Sholl intersection schematic of inset marked by dashed red box. Sholl intersection schematic depicts 10 µm steps from soma shown as concentric rings (red) with intersections marked with open dots. Scale bar, 50 µm. **(B-D)** Sholl analysis performed on day 35 FENIB KI neurons. The graphs depict mean ± SEM of n = 9-18 biological replicates (WT = 9, HET. KI = 10, HOM. KI = 10, CORR = 18). **(B)** Quantification of the number of dendritic intersections from 0-200 µm radial distances from the soma of day 35 FENIB KI neurons. (* are used to depict statistical significance between WT and comparison groups. # are used to depict statistical significance between CORR and comparison groups. *p<0.05, #p<0.05, ##p<0.01, ####p<0.0001). All comparisons left unmarked are statistically non-significant. **(C)** Quantification of the area under the curve for day 35 FENIB KI neurons. All comparisons left unmarked are statistically non-significant. **(D)** Quantification of total dendrite length for day 35 FENIB KI neurons. All comparisons left unmarked are statistically non-significant. **(E-G)** Sholl analysis performed on day 42 FENIB KI neurons. The graphs depict mean ± SEM of n = 15-17 biological replicates (WT = 15, HET. KI = 16, HOM. KI = 16, CORR = 17). **(E)** Quantification of the number of intersections from 0-200 µm radial distances from soma of day 42 FENIB KI neurons. (* are used to depict statistical significance between WT and comparison groups. # are used to depict statistical significance between CORR and comparison groups. *p<0.05, **p<0.01, #p<0.05, ##p<0.01). All comparisons left unmarked are statistically non-significant. **(F)** Quantification of area under the curve for day 42 FENIB KI neurons. (*p<0.05, **p<0.01). All comparisons left unmarked are statistically non-significant. **(G)** Quantification of total dendrite length for fay 42 FENIB KI neurons. (*p<0.05). All comparisons left unmarked are statistically non-significant. WT= wild-type, HET. KI = heterozygous knock-in, HOM. KI = homozygous knock-in, CORR = ABE corrected.

We first sought to assess differences between FENIB KI iNs at d35 **(Fig. 6B-D)**. We observed no significant differences in dendritic intersections between WT, heterozygous KI, homozygous KI, and ABE corrected neurons at d35 for any radial distance assessed. Intersection numbers peaked at 20 µm for WT (9.6 dendrites), heterozygous KI (9 dendrites), and ABE corrected (7.3 dendrites) and at 30 µm for homozygous KI (8.6 dendrites) (**Fig. 6B**). Using the number of intersections at radial distances, we calculated area under the curve (AUC) to determine overall dendritic arborization (**Fig. 6C**). Although WT and ABE corrected neurons appeared to have slightly greater AUC than both KI groups, statistical testing reveal no significant difference (WT: 819 arbitrary units (AU), heterozygous KI: 752 AU, homozygous KI: 620 AU, ABE corrected: 853 AU, p>0.05). Similarly, we quantified total dendritic length and found no statistically significant differences (WT: 1022.2 µm, heterozygous KI: 938.4 µm, homozygous KI: 864.4 µm, ABE corrected: 1177.5 µm, p>0.05) (**Fig. 6D**). We next sought to test differences in number of intersections at 10 µm increments from the soma across all groups using a two-way ANOVA with Bonferroni correction. We identified statistically significant differences in dendrite length in ABE corrected neurons compared to heterozygous KI and homozygous KI neurons at 70-130 µm radial distances from the soma (70 µm: ABE corrected: 6.2 intersections, homozygous KI: 3.6 intersections, p<0.05, 80 µm: ABE corrected: 5.8 intersections, homozygous KI: 2.9 intersections, p<0.01, 90 µm: ABE corrected: 5.9 intersections, homozygous KI: 2.7 intersections, p<0.01, heterozygous KI: 3.5, p<0.05, 100 µm: ABE corrected: 6.6 intersections, homozygous KI: 2.3, p<0.0001, heterozygous KI: 3.6, p<0.01, 110 µm: ABE corrected: 5.6 intersection, homozygous KI: 2.3 intersections, p<0.01, 120 µm: ABE corrected: 5.1 intersections, homozygous KI: 2.2 intersections, p<0.01, 130 µm: ABE corrected: 5.1 intersections, homozygous KI: 2 intersections, p<0.01). WT and ABE corrected neurons remained largely similar, with one statistically significant difference identified at 100 µm (ABE corrected: 6.6 intersections, WT: 3.6 intersections, p<0.05).

As neuroserpin functions in late-stage neuronal maturation, we allowed further neuronal development and assessed dendritic morphology again at d42 (**Fig. 6E-G)**. Intersection number peaked at 30 µm for WT (11.6 dendrites), homozygous KI (7.9 dendrites), and ABE corrected neurons (11.8 dendrites), and at 20 µm for heterozygous KI (8.4 dendrites) (**Fig. 6E**). When we compared AUC for d42 neurons, we identified a significant impairment in overall dendritic arborization in heterozygous and homozygous KI neurons compared to WT (WT: 1293 AU, heterozygous KI: 822.5 AU, homozygous KI: 887.8 AU, p<0.01 for WT vs. heterozygous KI, p<0.05 for WT vs. homozygous KI) (**Fig. 6F**). Comparison of heterozygous and homozygous KI to ABE corrected neurons was trending but statistically non-significantly (ABE corrected: 1174 AU, p>0.05). When we compared AUC between WT and ABE corrected neurons, we identified no differences in overall dendritic arborization (p>0.05), suggesting that ABE correction results in neurons with dendritic arborization phenotype in-between WT and KI variant neurons. We identified a similar impairment when we assessed total dendritic length in d42 neurons (**Fig. 6G**). Heterozygous and homozygous KI neurons had reduced total dendritic length compared to WT control neurons (WT: 1704.7 µm, heterozygous KI: 1086.4 µm, homozygous KI µm: 1093.8 µm, p<0.05). Once again, ABE corrected neurons were statistically non-significant from either WT or KI neurons (ABE corrected: 1536 µm, p>0.05). Next, we ran a two-way ANOVA with Bonferroni correction to test differences in number of intersections at 10 µm radial distances from the soma across WT, heterozygous KI, homozygous KI, and ABE corrected iNs. Similar to d35, we identified statistically significant differences between both WT and heterozygous and homozygous KI neurons, and ABE corrected and heterozygous and homozygous KI neurons. These differences occur at 20-40 and 60-90 µm radial distances from soma for WT neurons (20 µm: WT: 11.4 intersections, homozygous KI: 7.0 intersections, p<0.01, 30 µm: WT: 11.6 intersections, heterozygous KI: 7.8 intersections, p<0.05, homozygous KI: 7.9 intersections, p<0.05, 40 µm: WT: 10.7 intersections, homozygous KI: 7.4 intersections, p<0.05, 60 µm: WT:10.5 intersections, heterozygous KI: 6.5 intersections, p<0.01, homozygous KI: 7.1 intersections, p<0.05, 70 µm: WT: 10.2 intersections, heterozygous KI: 6.1 intersections, p<0.01, homozygous KI: 5.9 intersections, p<0.01, 80 µm: WT: 9.1 intersections, heterozygous KI: 5.1 intersections, p<0.01, homozygous KI: 5.4 intersections, p<0.05, 90 µm: WT: 8.8 intersections, heterozygous KI: 4.3 intersections, p<0.01, homozygous KI: 5.3 intersections, p<0.05) and 20- 40 µm radial distances for ABE corrected neurons (20 µm: ABE corrected: 10.4 intersections, homozygous KI: 7.0 intersections, p<0.05, 30 µm: ABE corrected: 11.8 intersections, heterozygous KI: 7.8 intersections, p<0.001, homozygous KI: 7.9 intersections, p<0.01, 40 µm: ABE corrected: 11.1 intersections, heterozygous KI: 7.9 intersections, p<0.05, homozygous KI:7.4 intersections, p<0.05). These findings support a developmental impairment arising from the mutant neuroserpin activity in addition to the toxic gain-of-function of aggregating mutant neuroserpin.

### eVLP-mediated delivery of ABEs can efficiently correct the FENIB variant

Delivery of gene editors to target tissues is an outstanding challenge for therapeutic genome editing.^36,70^ We decided to test eVLPs to target neurons due to the high cell-targeting efficiency and subsequent editing, and simplicity in reconfiguring eVLP constructs. We produced eVLPs by transfecting HEK293T cells with plasmids encoding VSV-G glycoprotein, MMLV-gag-pol, and MMLV fused NG-ABE8e ribonucleoprotein, henceforth called the eVLP plasmids (**Fig. 7A**). We first tested eVLP editing on target loci with documented high editing; *HEK2* and *HEK3* (**Fig. 7B and 7C**).^39^ To generate eVLPs, we transfected HEK293T cells with the eVLP plasmids, and collected the eVLP containing media after 72 hours.^39^ We concentrated the supernatant 100-fold using PEG-IT, a polyethylene glycol (PEG) based reagent. Next, we treated HEK293T cells with 100-fold concentrated eVLPs and sequenced the target loci 72 hours after treatment. We observed robust editing at both the *HEK2* and *HEK3* loci, with the greatest editing at the A7 position (88%) and A6 position (64%), respectively.

**Figure 7.**
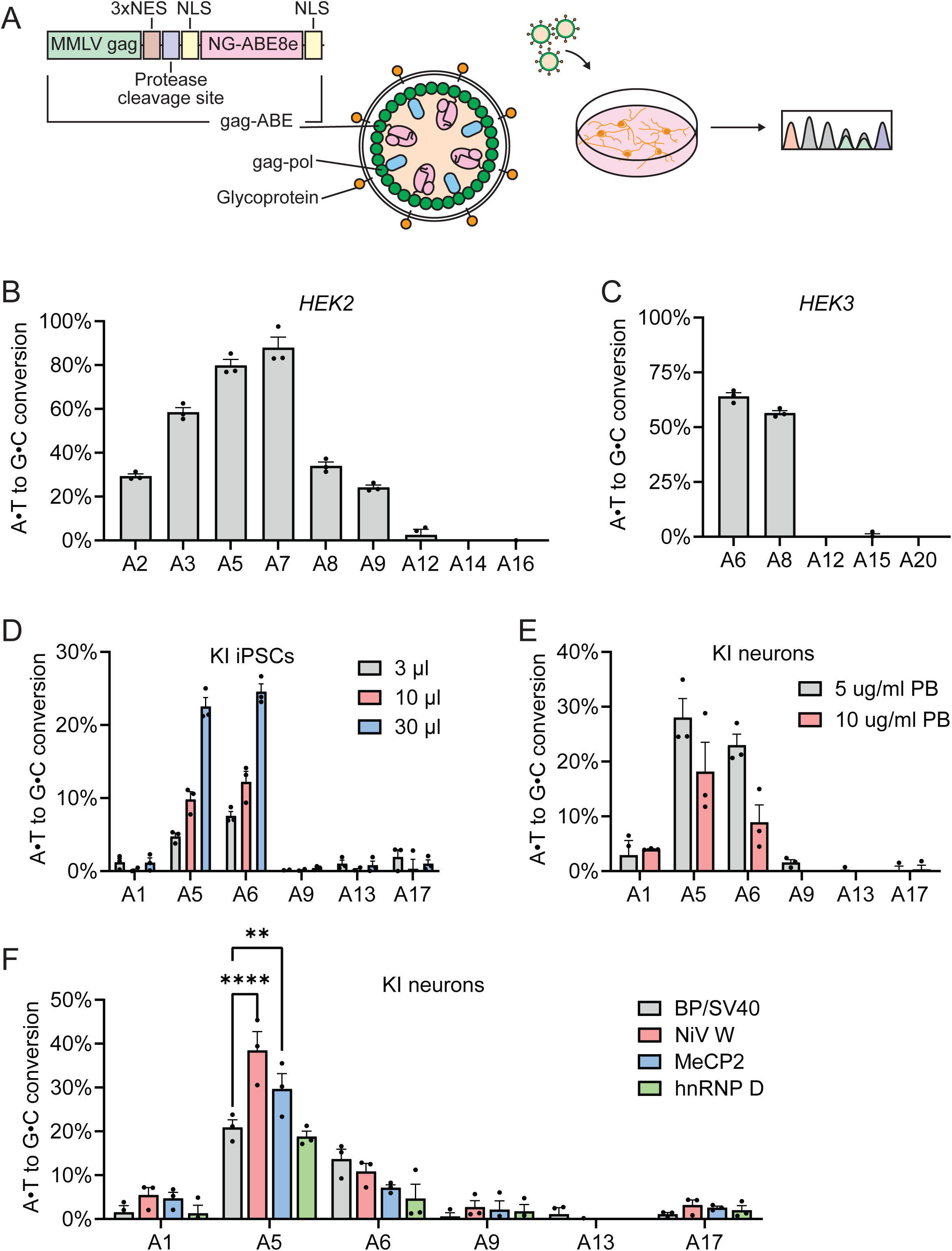
eVLP editing in iPSC and iPSC-derived neurons. **(A)** MMLV-gag fused NG-ABE8e for eVLPs (top), structure of ABE containing eVLP (middle), and depiction of editing and sequencing (bottom). MMLV-gag (green) fused NG-ABE8e (pink) with nuclear localization signals (brown), nuclear localization signal (yellow), and protease cleavage site (blue). eVLPs generated using MMLV-gag-pol (green and blue), VSV-G or FuG-B2 glycoprotein (orange), and MMLV-gag-NG-ABE8e (green and pink) (middle). Diagram of eVLP treated cells and subsequent editing at target locus. **(B)** Quantification of eVLP-ABE editing at the HEK2 locus in HEK293T cells. The graphs depict mean ± SEM of n = 3 biological replicates. **(C)** Quantification of eVLP-ABE editing at the HEK3 locus in HEK293T cells. The graphs depict mean ± SEM of n = 3 biological replicates. **(D)** Quantification of eVLP-ABE correction of FENIB mutation in *SERPINI1* c.1175 homozygous KI iPSCs. The graphs depict mean ± SEM of n = 3 biological replicates. **(E)** Quantification of eVLP ABE editing in day 7 iPSC-derived neurons using 5 µg/ml polybrene (PB) with BP/SV40 NLS (grey) or NiV NLS (red) or 10 µg/ml PB with BP/SV40 NLS (blue) or NiVW NLS (green). The graphs depict mean ± SEM of n = 3 biological replicates. **(F)** Quantification of eVLP-ABE correction of FENIB mutation in d7 iPSC-derived neurons using neuron-specific nuclear localization signals NiV W (red), MeCP2 (blue), hnRNP D (green) or BP/SV40 (grey) control. The graphs depict mean ± SEM of n = 3 biological replicates. (****p<0.0001, **p<0.001.) All comparisons left unmarked are statistically non-significant.

After confirming the capacity of eVLPs to deliver ABE and edit target loci in HEK293T cells, we sought to test eVLP editing in *SERPINI1* KI iPSCs. We treated homozygous KI iPSCs with eVLP containing supernatant in Stemflex iPSC media at ratios of 1:10, 1:3, and 1:1 (**Fig. S7B**). After 72 hours, we collected DNA from the treated cells and Sanger sequenced the samples at the *SERPINI1* c.1175 G>A locus. Without concentration or selection, we observed >10 % editing at the A5 and A6 positions in our 1:1 ratio of eVLP containing supernatant to Stemflex media when using EditR Sanger sequencing analysis. We next generated concentrated eVLPs and treated homozygous KI iPSCs and sequenced the target locus 72 hours after treatment without selection (**Fig. 7D**). eVLP editing using EditR revealed a dosage effect, with the 30 µl high dose editing resulting in 22.5% correction to the wild-type sequence at the A5 position and 25% correction at the A6 position.

We assessed neural editing by treating homozygous KI neurons with optimized eVLP conditions. Concentrated eVLPs were added with neurotrophic factors, ARAC, and doxycycline at d7 of neural induction. Treated neurons were collected without selection and Sanger sequenced at the target locus after 72 hours (**Fig. S7C**). We observed a dosage effect in our homozygous KI iNs. However, our highest dose, 30ul, only reached 2% at the A5 and 1.5% editing at the A6 position, necessitating further optimization.

We tested a modified rabies virus glycoprotein, FuG-B2, to increase eVLP uptake through enhancing neuronal specificity and selectivity.^39,71^ Homozygous KI neurons treated with concentrated eVLPs enveloped with FuG-B2 glycoprotein increased editing to 1% at the A5 and 3% at the A6 positions (**Fig. S7D**). To optimize neuron specific nuclear localization, we generated NG-ABE8e flanked by Nipah virus W protein (NiV W), methyl CpG binding protein 2 (MeCP2), and heterogenous nuclear ribonucleoprotein D (hnRNP D) NLSs.^72^ We produced each NLS variant in eVLPs with both VSV-G and FuG-B2 glycoproteins. For VSV-G eVLPs, NiV W NLS (1.7%) resulted in the highest correction at the A5 position, followed by MeCP2 NLS (0.6%), BP/SV40 NLS (0.5%), and hnRNPD NLS (0%) (**Fig. S7E**). FuG-B2 eVLP results were similar with NiV W NLS (3.2%) resulting in the highest correction efficiency, followed by BP/SV40 NLS (2.9%), MeCP2 NLS (0.8%), and hnRNPD NLS (0%) (**Fig. S7F**). The difference in eVLP editing efficiencies between HEK293T cells and iPSCs, and iPSC-derived neurons suggested an issue with eVLP uptake.

We hypothesized that eVLP-cell interactions were insufficient for eVLP uptake and decided to test the addition of polybrene (**Fig. 7E**). Polybrene is a cationic polymer which facilitates receptor-independent virus adsorption.^73^ Both 5 µg/ml and 10 µg/ml polybrene increased eVLP editing by 10-fold in VSV-G eVLPs. eVLP editing in homozygous KI neurons with VSV-G glycoprotein with 5 µg/ml polybrene had the highest editing efficiency at the A5 position (28%) while 10 µg/ml polybrene followed closely (18%).

We assessed eVLPs with neuron specific NLS with 5 µg/ml polybrene to determine optimal neuron editing. We treated homozygous KI neurons with VSV-G glycoprotein eVLPs and either BP/SV40 or neuron-specific NLS (**Fig. 7F**). Homozygous KI neurons treated with eVLPs with NiV W NLS (38.5%, p<0.0001) had the highest editing efficiency, significantly greater than BP/SV40 (20.9%). We also found that MeCP2 NLS (29.7%, p<0.01) had statistically increased editing at the A5 position. Editing between hnRNP D NLS and BP/SV40 was statistically non-significant (18.8% p=0.76). In addition to differences in editing efficiency, we found that neuron specific NLS eVLPs had different editing profiles than standard BP/SV40. There was low but detectable editing at all A positions along the gRNA. BP/SV40 had the broadest editing window with A5 (20.9%) A6 (13.7%). The neuronal NLS editors had a higher ratio of desired A5 to bystander A6 editing, with NiV W 38.5%:10.9%, MeCP2 29.7%:7.2%, and hnRNP D at 18.8%:4.7%.

We next assessed eVLP editing of MUT NS-GFP neurons (**Fig. S7G**). VSV-G with BP/SV40 NLS (13.4%) had greater editing efficiency at the A5 position. Unlike homozygous KI neurons, eVLP editing of the MUT NS-GFP loci by NiV W (4.8% p<0.0001), MeCP2 (0.4%, p<0.0001), and hnRNP D (0.16% p<0.0001) were statistically less efficient than BP/SV40. Thus, neuron specific NLS eVLPs increase editing efficiency in KI neurons and increased the desired A5 to bystander A6 editing ratio, but not in the MUT NS-GFP neurons.

## Discussion

We aimed to generate patient-specific models of protein conformational disease to assess whether ABEs can efficiently correct the pathogenic variant, and if preventing novel mutant protein expression would facilitate clearance of neuroserpin aggregates. Our results support that CRISPR/Cas-mediated genome editing systems can model and eliminate aggregates or prevent aggregation progression in this rare patient-specific variant of brain disease using HEK293T cells, iPSCs, and neuron models.

We hypothesized that disease modifying treatments for FENIB would require resolving NS aggregation, similar to potential therapies for Alzheimer’s and Huntington’s diseases which target aggregate-prone protein precursors.^16,19^ Doxycycline withdrawal experiments demonstrated that early prevention of mutant protein expression facilitates complete clearance of NS-GFP aggregates. Whether clearance of aggregates resulted from reduced aggregate burden or because neurons were younger remains unclear. The average number of aggregates per cell and size of aggregates were similar in d10 neurons and d21 neurons, suggesting that the difference in aggregate clearance reflected neuron age, not aggregate burden. In more mature neurons, we were unable to achieve complete clearance, but prevented further NS-GFP aggregation. Coupling prevention of novel protein production with methods to promote protein turnover may facilitate aggregate clearance. Embelin, a small molecule derived from the Japanese Ardisia (*Ardisia japonica*) herb, prevented *in vitro* NS aggregation and dissolved aggregates.^74,75^ ABE correction of the pathogenic variant to prevent new aggregation combined with embelin disruption of existing aggregates is a potential effective therapy.

Research of conformational neurodegenerative diseases is often limited by experimental timescale where aggregate burden requires aging over years or decades for accumulation and neurotoxicity.^76,77^ Overexpression of NS overcomes low levels and reduced overall duration of endogenous NS protein expression, and has been used to successfully study FENIB and NS aggregation both *in vitro* and *in vivo*.^5,14,46,49,63,63,78–81^ However, overexpression of NS-GFP to study aggregation obscures the potential phenotypes which may arise given patients’ heterozygosity. We complemented NS-GFP with heterozygous and homozygous KI neurons and identified a novel developmental deficit in dendritic arborization and total dendritic length. Dendritic spines are directly involved in receiving excitatory synaptic inputs in excitatory cortical neurons. Impaired dendritic spines are a known abnormality in epilepsy, but it remains unclear whether this is a cause or result of epilepsy.^82^ Dendritic pathologies have been also identified in neurological disorders with epilepsy including Rett and Down syndrome.^83,84^ Approximately 90% of patients with Rett syndrome have epilepsy. Knockdown of endogenous *Mecp2*, the gene implicated in Rett syndrome, using shRNAs reduced the number of dendritic nodes, total dendritic length, and dendritic intersections in a neuronal model.^85^ Further analysis of dendritic spines in FENIB will be illustrative to assess whether heterozygous and homozygous KI neurons suffer impairments to spine morphology in addition to dendritic arbor.

Whole-cell patch clamp electrophysiology revealed no differences between WT and heterozygous or homozygous KI neurons, suggesting that partial or complete loss of wild-type NS protein does not directly cause seizures. Because our model generates cortical excitatory neurons, it remains possible that other neuronal subtypes, specifically inhibitory neurons, could be affected in FENIB. Additionally, our *in vitro* cultures may not sufficiently model the interactions between NS and tissue-type plasminogen activator (tPA).^86–88^ tPA enhances brain N-methyl-D-aspartate receptor (NMDAR) activity through a mechanism that has yet to be completely understood.^89,90^ NS protects against NMDAR excitotoxicity *in vivo* and *in vitro* by limiting tPA’s pro-excitotoxic effects.^91^ Likely, further *in vivo* modeling of aged FENIB neurons will be required to assess whether the NS-tPA and NMDAR interactions are responsible for the epileptic phenotype.

Delivery of genome editors larger than 5kb remains a challenge. This is due to the genomic packaging limitations of AAVs. eVLPs with neuron-specific NLSs improved editing efficiency of *in vitro* cortical excitatory neurons with the additional benefit of narrowing the on-target: bystander editing ratio. The delivery method of genome editors affects the duration of Cas9 expression. Plasmids-based methods results in days, AAV-based methods in months, and lentivirus-based methods in lifelong expression. eVLPs deliver genome editors as ribonucleoprotein, resulting in the turnover of most protein after 24 hours. ^92^ We hypothesize that due to the minimal window of expression, eVLP mediated ABE delivery requires faster nuclear import observed when using neuron-specific NLSs. ^72^ For this reason, we envision that neuron-specific NLS’s will improve editing with both AAV and lipid-nanoparticle RNPs, as both require NLSs for nuclear entry.

In summary, our CRISPR/Cas genome editors effectively modeled and corrected the severe patient-specific FENIB variant. Our modeling strategy uses complementary cellular systems to investigate partial or complete loss of wild-type NS and toxic gain-of-function present in FENIB. Heterozygous and homozygous KI neurons revealed developmental deficits resulting in impaired dendritic morphology. Early termination of MUT NS-GFP expression can completely clear aggregates, while late termination prevented further NS-GFP aggregation. Treatment of MUT NS-GFP cells with ABE facilitated NS-GFP clearance. Finally, we increased editing efficiency in neurons using NiV W NLS peptide to shuttle NG-ABE8e into neuronal nuclei.

## Materials and Methods iPSC culture

Induced pluripotent stem cells (iPSC) were cultured in Stemflex media (Gibco; Grand Island, NY) supplemented with 1% antibiotic-antimycotic (Gibco; Grand Island, NY) with daily media changes. Cells were split and passaged at 70% confluency, approximately every 3-4 days, using 0.5mM EDTA (Invitrogen; Waltham, MA) in DPBS (Gibco; Grand Island, NY) or Accutase (Innovative Cell Technologies; San Diego, CA) and plated onto geltrex (Gibco; Grand Island, NY) coated tissue culture treated wells. When passing, iPSCs were plated in Stemflex media with 1% antibiotic-antimycotic supplemented with 10uM ROCK-inhibitor (LC Laboratories; Woburn, MA). Cell media was changed with Stemflex media without ROCK-inhibitor the following day. Cells were maintained at 37°C with 5.0% CO_2_.

## Neural induction

iPSC-derived neurons were generated through doxycycline (Sigma Aldrich; Saint Louis, MO) induced expression of Ngn2.^31^ Briefly, iPSCs were detached and passaged onto geltrex coated plates, and transduced with lentivirus expressing ubiquitin-C promoter expressed reverse tetracycline transactivator, or tetracycline response element On promoter expressed mouse Neurogenin2-T2A-puromycin (*Ngn2*) resistance in Stemflex media with 10uM ROCK-inhibitor. On day 1, culturing media was changed to iN media consisting of Neurobasal medium (Gibco; Grand Island, NY) with 1X B-27 (Gibco; Grand Island, NY) and further supplemented with 10 µM ROCK-inhibitor and doxycycline to induce *Ngn2* expression. Culturing media was replaced every day with iN media until day 6, with doxycycline induction from days 1-7, and puromycin (Invivogen; San Diego, CA) selection from days 2-5. On day 6, cells were detached and replated onto glia isolated from p3 mouse pups in iN media supplemented with 5% FBS (GeminiBio; West Sacramento, CA) on either geltrex coated glass coverslips for immunocytochemical labeling and patch clamp electrophysiology or plastic tissue culture plates for RNA and protein processing. Beginning d7, culturing media was replaced every 2-3 days with iN media supplemented with BDNF (STEMCELL Technologies; Vancouver, Canada) GDNF (STEMCELL Technologies, Vancouver, Canada), NT-3 (STEMCELL Technologies; Vancouver, Canada), and ARAC (Selleckchem; Houston, TX). Cells were maintained at 37°C with 5.0% CO_2_.

## HEK293T cell culture

HEK293T cells were cultured in media consisting of DMEM (Gibco; Grand Island, NY) supplemented with 10% fetal bovine serum (FBS) and 1% antibiotic-antimycotic with media changes every 2-3 days. Cells were passaged every 5-7 days or when 70% confluent using 0.25% trypsin/EDTA (Gibco; Grand Island, NY). Cells were maintained at 37°C with 5.0% CO_2_.

## Plasmids

The plasmid pSpCas9(BB)-2A-GFP (PX458) was a gift from Feng Zhang (#48138, Addgene), the plasmid pTet-O-Ngn2-puro was a gift from Marius Wernig (#52047, Addgene), the plasmid FUW-M2rtTA was a gift from Rudolf Jaenisch (#20342, Addgene), the plasmids pCMV_ABEmax (#112095, Addgene), ABE8e (#138489, Addgene), and pCMV-MMLVgag-3xNES-ABE8e-NG (#181754, Addgene) were gifts from David Liu, the plasmids SECURE miniABEmax-V82G (pJUL1828) (#131313, Addgene) and SECURE miniABEmax-K20A/R21A (pJUL1774) (#131312, Addgene) were gifts from Keith Joung, the plasmid pBS-CMV-gagpol was a gift from Patrick Salmon (#35614, Addgene), the plasmid pCMV-VSV-G was a gift from Bob Weinberg (#8454, Addgene), the plasmid pCAGGS-FuG-B2 was a gift from Kazuto Kobayashi (#67513, Addgene), the plasmids cTNT-BLAST-T2A-eGFP-RP, psPAX2, and pMD2G were gifts from Dr. Lei Bu, and the plasmid pBSU6_FE_Scaffold_RSV_GFP was a gift from Dr. Dirk Grimm.

The plasmid SpCas9-NG was generated by introducing the mutations L1111R, D1135V, G1218R, E1219F, A1322R, R1335V, T1337R into the pSpCas9(BB)-2A-GFP (PX458) plasmid using primer directed mutagenesis and Gibson assembly. ^41^ gRNAs targeting *SERPINI1* c.1175 G>A were cloned into the plasmid pBSU6_FE_Scaffold_RSV_GFP. The C-terminal portion of SpCas9-NG fused to 2A-GFP expressing fragment was amplified from the SpCas9-NG plasmid and cloned into the ABEmax, ABE8e, V82G, and K20A/R21A ABE plasmids using Gibson assembly. pTet-O-WT-NS-GFP-BSD and pTet-O-MUT-NS-GFP-BSD plasmids were cloned into pTet-O-Ngn2-puro plasmids. Wild-type (WT) and mutant (MUT) *SERPINI1* cDNA were amplified from cDNA synthesized from wild-type iPSC-derived neurons and homozygous KI iPSC-derived neurons expressing *SERPINI1* transcripts, respectively. The GFP expressing fragment was amplified from pSpCas9(BB)-2A-GFP. A 3X glycine-glycine-glycine-serine linker sequence was added between the *SERPINI1* and GFP sequences by primer directed mutagenesis. The 2A- BSD expressing fragment was amplified from cTNT-BLAST-T2A-eGFP-RP. The *SERPINI1*, GFP, and 2A-BSD fragments were cloned into the pTet-O-Ngn2-puro plasmid using Gibson assembly. CMV-MMLVgag-3xNES-NiVW-ABE8e-NG, pCMV-MMLVgag-3xNES-MeCP2-ABE8e-NG, and pCMV-MMLVgag-3xNES-hnRNPD-ABE8e-NG plasmids were generated by replacing the SV40 and BP NLS sequences in the pCMV-MMLVgag-3xNES-ABE8e-NG plasmid with neuron-specific NLS sequences using primer directed mutagenesis and Gibson assembly. All Gibson assembly primers are listed in Table 1, all plasmid sequences are listed in Table 2, all plasmid sequencing primers are listed in Table 3.

Plasmids were transformed into DH5A (Invitrogen; Waltham, MA) or Stbl3 (Invitrogen; Waltham, MA) bacteria, and amplified using Nucleobond Xtra Maxi plasmid purification kit (Macherey-Nagel; Allentown, PA). Plasmid sequences were verified using Sanger sequencing or Plasmid-EZ (Genewiz; Waltham, MA).

## Transfections

Transfections were performed using nucleofections, lipofectamine, or polyethylenimine. Nucleofections used 4D nucleofector system with P3 or SF reagent (Lonza; Morris Township, NJ) for iPSCs and HEK293T cells respectively, per manufacturer’s guidelines. For generating the knock-in model, cells were nucleofected with 5 µg px458 or NG-px458 plasmid with locus specific gRNA and 5 µl of 10 µM ssODN (Integrated DNA technologies; Coralville, IA). Lipofection was performed using lipofectamine 2000 (Invitrogen; Waltham, MA) as reverse transfections, per manufacturer’s guidelines. PEI (Sigma Aldrich; Saint Louis, MO) transfections were performed as reverse transfections, using a 1:5 ratio of plasmid to PEI in Opti-MEM (Gibco; Grand Island, NY) All gRNA sequences are listed in Table 4.

## eVLP and lentivirus preparation

Lentivirus and eVLP vectors were produced by reverse transfection of 1.2 x 10^7^ HEK293T cells using PEI in Opti-MEM. To generate lentivirus, cells were reverse transfected with psPAX2 (4.7 µg), pMD2.G (1.6 µg), and expression plasmids (6.3 µg) per 10 cm dish. To generate eVLPs, cells were reverse transfected with VSV-G or FuG-B2 (400 ng), MMLVgag-pro-pol (3,375 ng), MMLVgag-3XNES-NG-ABE8e or neuron-specific NLS variants (1,125 ng), and sgRNA (4,400 ng) plasmids per 10 cm dish. 24-hours post-transfection, culturing media was replaced with DMEM+/+. 72-hours post-transfection, vector containing media was harvested and centrifuged for 10 min at 500 RCF to pellet cells. Vector containing supernatant was filtered using a 0.45 µm PVDF syringe filter (Corning; Corning, NY). Vector supernatant was concentrated 100-fold using Lenti-X concentrator (Takara Bio; Shiga, Japan) or PEG-IT virus precipitation solution (System Biosciences; Palo Alto, CA) per manufacturer’s instructions in cold 1 X DPBS. Concentrated vectors were either immediately used or stored at -80°C avoiding multiple freeze-thaws.

## Flow cytometry

HEK293T cells and iPSCs were dissociated into a single cell suspension using 0.25% trypsin/EDTA or Accutase, respectively, and sorted 72-hours post-transfection using either a MoFlo-XDP or SY3200 cell sorter. Cells were gated via doublet discrimination, followed by live/dead viability based on DAPI staining. Cells were sorted by GFP or GFP and RFP signal based on the experiment. Cells were sorted using a 100 µm nozzle. To generate stable clones, cells were single-cell sorted into 96-well plates into DMEM +/+ for HEK293T cells or Stemflex supplemented with 1X CloneR (STEMCELL Technologies; Vancouver, Canada) for iPSCs.

## DNA extraction

DNA extraction was performed on a minimum of 10,000 cells. Cells were processed with DirectPCR lysis-reagent cell (Viagen; Los Angeles, CA) to extract crude DNA. Crude DNA was directly used for PCR.

## PCR

GoTaq PCR (Promega; Madison, WI) was used to generate amplicons using sequence-specific primers (Integrated DNA technologies; Coralville, IA) in the Mastercycler nexus X2 thermal cycler (Eppendorf; Hamburg, Germany). Primer sequences can be found in Table 5.

## T7E1, Synthego ICE, and EditR

T7 endonuclease I (T7E1) assay was performed by reannealing raw PCR product from GoTaq PCR amplification followed by incubation of reannealed product with T7E1 for digestion of mismatched paired strands. Synthego ICE analysis was performed by comparing Sanger sequencing .ab1 files per application instructions.^44^ Briefly, .ab1 Sanger sequencing traces from edited samples are provided along with unedited control traces, sgRNA sequences, and HDR KI template sequences. These files are used to compare the edited sequence against the unedited control and determine the percentage of signal corresponding to the intended KI sequence, nonspecific indels, and unedited wild-type sequence. EditR analysis was performed by calculating the difference in base calls at the target locus between ABE treated and unedited control cells as determined by EditR software (1.0.10).^62^ Sanger sequencing trace files were uploaded in combination with gRNA sequence, with P-value cutoff at the default value, 0.01. 5’ start and 3’ ends were determined empirically based off Sanger sequencing trace quality for each locus tested as follows: *SERPINI1* 5’ start: 50bp, 3’ end: 280 bp, *HEK2* 5’ cut start: 50 bp, 3’ end: 280 bp, *HEK3* 5’ start: 50 bp, 3’ end: 550 bp.

## Immunocytochemistry

Cells were plated on geltrex coated 12 mm No.1 coverslips (Chemglass Life Sciences; Vineland, NJ) in a 24-well plate. At experimental endpoints, cells were washed with DPBS to remove residual media, then fixed in 4% PFA (Electron Microscopy Sciences; Hatfield, PA) in PBS at room temperature. Samples were subsequently permeabilized using 0.1% triton-x 100 (Sigma Aldrich; Saint Louis, MO) in PBS, washed three times with PBS, then blocked using 10% either donkey serum (Abcam; Waltham, MA), 0.25% triton-x 100 in PBS, and washed three times with PBS. Cells were incubated with primary antibodies overnight at 4°C. The following antibodies were used for immunocytochemistry: Rabbit anti-neurofilament 200 (1:1000, Sigma Aldrich, N4142-25ul), rabbit anti-neuroserpin (1:200, Abcam, ab33077), rabbit anti-synapsin 1 (1:1000, Sigma Aldrich, AB1543), mouse anti-NeuN (1:500, Abcam, ab104224), mouse anti- MAP2 (1:500, Sigma Aldrich, MAB3418), rabbit anti-synapsin 1 (1:1000, Invitrogen, A-6442), mouse anti-SSEA4 (1:250, Invitrogen, MC-813-70), rabbit anti-SOX2 (1:250, Invitrogen, PA1- 094), mouse anti-GFP (1:1000, MBL Life Science, M048-3), rabbit anti-neuroserpin (1:1000, a gift from Dr. M. Elena Miranda Banos, purified polyclonal antibody) and mouse anti-neuroserpin (1:1000, a gift from Dr. M. Elena Miranda Banos, 1A10). ^63^ The following day, cells were washed three times with PBS and incubated with secondary antibodies in 10% donkey serum in PBS for 1 hour at room temperature. The following secondary antibodies were used: donkey anti-mouse IgG (H+L) highly cross-adsorbed secondary antibody Alexa Fluor 488 (1:250, Invitrogen, A-21202), donkey anti-mouse IgG (H+L) highly cross-adsorbed secondary antibody Alexa Fluor 594 (1:250, Invitrogen, A-21203), donkey anti-mouse IgG (H+L) highly cross-adsorbed secondary antibody Alexa Fluor 647 (1:250, Invitrogen, A-31571), donkey anti-rabbit IgG (H+L) highly cross-adsorbed secondary antibody Alexa Fluor 488 (1:250, Invitrogen, A-21206), donkey anti-rabbit IgG (H+L) highly cross-adsorbed secondary antibody Alexa Fluor 594 (1:250, Invitrogen, A-21207). Nuclei were stained with DAPI (Invitrogen; Waltham, MA), then stained slides were mounted with fluorescence mounting medium (Dako; Glostrup, Denmark). Cells were imaged with a Zeiss-880 inverted confocal microscope with a magnification of 40X N.A. 1.1, 1 Airy unit pinhole, half-maximal optimized Z-stack increments with 2x2 tiled images.

## Western Blot

Cells were lysed using a lysis buffer containing 20 mM Tris-HCl, pH 7.5, 150 mM NaCl, cOmplete mini EDTA-free protease inhibitor cocktail (Roche; Indianapolis, IN), PhosSTOP phosphatase inhibitor cocktail (Roche; Indianapolis, IN), 1% Triton-X 100, in PBS. Samples were incubated on ice for 30 minutes and vortexed every 10 minutes. Samples were centrifuged at 20,000 RCF for 30 minutes at 4°C to remove insoluble material.

Protein concentration was determined using the Pierce BCA kit (Thermo Fisher Scientific; Waltham, MA). 25 µg of whole cell lysate was loaded into a 7.5% mini-protein precast protein gel (Bio-Rad; Hercules, CA). Lysate for SDS-PAGE was prepared in 4X sample buffer with 2- mercaptoethanol (Sigma Aldrich; Saint Louis, MO) and heated at 98°C for 5 minutes. Lysate for native-PAGE was prepared with BlueJuice Gel loading buffer (Thermo Fisher Scientific; Waltham, MA). Gels were run at 180V for 1 hour at 4°C in either SDS-PAGE (25 mM Tris, 192 mM glycine, 1% SDS in diH2O) or native-PAGE running buffer (25 mM Tris, 192 mM glycine in diH2O) to separate proteins. Gels were electrophoretically transferred onto Immun-Blot PVDF membrane (Bio-Rad; Hercules, CA) overnight in transfer buffer (25 mM Tris, 190 mM glycine, 0.1% SDS, 20% methanol in diH2O) at a constant 20mA. Membranes were washed with TBST (20 mM Tris, 150 mM NaCl, 0.1% tween 20 in diH2O) then blocked in 5% nonfat dry milk in TBST. Membranes were rotationally incubated with diluted primary antibodies in 5% nonfat dry milk in TBST at 4°C overnight. The following antibodies were used for immunoblotting: rabbit anti-neuroserpin (1:200, Abcam, ab33077), mouse anti-vinculin (1:1000, Sigma Aldrich, V9131- 2ML), mouse anti-GFP (1:1000, MBL Life Science, M048-3). The following day, membranes were washed with TBST and incubated for 1 hour at room temperature with either goat anti- mouse IgG (H+L) secondary antibody conjugated to HRP (1:5000, Invitrogen, 31430) or goat anti-rabbit IgG secondary antibody conjugated to HRP (1:5000, Invitrogen, 31460) in TBST. Membranes were washed three times with TBST, and a final wash with TBS before applying Western Blotting Luminol Reagent (Santa Cruz Biotechnology; Dallas, TX) and imaging with a Bio Rad ChemiDoc imaging system.

### RNA extraction and cDNA synthesis

RNA was extracted using RNeasy mini kit (Qiagen; Hilden, Germany) with on-column DNase digestion (Qiagen; Hilden, Germany) per the manufacturer’s guidelines. cDNA was synthesized from RNA using the SuperScript III First-Strand cDNA Synthesis Kit (Thermo Fisher Scientific; Waltham, MA).

### Quantitative PCR

Quantitative PCR was performed on cDNA synthesized as mentioned above using ssoAdvanced Universal SYBR Green Supermix reagent (Bio-Rad; Hercules, CA). For each assay, 500 nM of forward and reverse primers, and 1X final concentration of SYBR green master mix was used. Assays were performed using a StepOne Real-time PCR system (Applied Biosystems; Waltham, MA) using the following settings: 30 seconds at 95°C, followed by 40 cycles of 10 seconds at 95°C and 60 seconds at 60°C, followed by a melt curve analysis of 40 cycles at 65°C -95°C increasing in 0.5°C increments at a rate of 5 seconds per step. All qPCR primers are listed in table 6.

### Next generation sequencing

RNA was extracted using RNeasy mini kit as mentioned above. Automated polyA selected stranded RNA-seq library prep was performed by the New York University Langone Health Genome Technology Center. Libraries were sequenced using an Illumina NovaSeq6000 next generation sequencer (Illumina Inc; San Diego, CA) as paired end runs. After quality control, samples were mapped using BBTools software, Sequence Expression Analyzer (Seal) (Joint Genome Institute; Berkeley, CA) against both mouse and human reference genomes to exclude reads from mouse glia.^93^ Reads were mapped using Spliced Transcripts Aligned to a Reference (STAR), and counted with featureCounts.^94,95^ Read counts were normalized as transcripts per million (TPM) and plotted as log_2_(TPM+1).

### Aggregate analysis and detection

Aggregate analysis was performed using ImageJ FIJI software. Confocal microscopy acquired images were projected as maximum projections of Z-stack images to create a two-dimensional image of aggregates. The NS-GFP channel was separated and thresholded for the brightest fluorescence expression. All images from one experiment were thresholded using the same values. Aggregates were counted using ImageJ “analyze particles” function. Particles of less than 0.1 microns were excluded to avoid including single pixel values in the analysis. The total number of particles in an image was used to determine aggregate count. The total area of the particles in an image was used to determine the aggregate area.

NS-GFP aggregate detection for single clone assay was performed using the Olympus CKX53 epifluorescence microscope (Evident Corporation, Tokyo, Japan). GFP fluorescence and brightfield images were acquired using cellSens imaging software (Evident Corporation, Tokyo,

Japan). All images were acquired under the same gain and exposure settings. Images were processed using ImageJ FIJI software. The NS-GFP channel was thresholded for the brightest fluorescence to detect NS-GFP aggregates. Images were categorized as containing aggregates if any particle greater than 0.1 micron was detected.

### Cell count

Confocal microscopy acquired images were used to identified and count cells using Cellpose v2.2.2 software.^96^ Images were initially processed in ImageJ to isolate the DAPI stained nuclei channel, and project using the sum projection option to produce an image of all cells within the field-of-view. The nuclei channels were processed in Cellpose set to nuclei mode. Cell diameter was automatically determined using the calibration function. Flow_threshold was set to 0.4, cellprob_threshold was set to 0.0, and stitch_threshold was set to 0.0. Image saturation was set to automatic. The resulting cell count was used to determine the total number of cells per field. To determine NeuN and DAPI double positive cells, images of NeuN and DAPI stained cells were processed using ImageJ. The NeuN channel was isolated and automatically thresholded to generate a binary mask. The mask was processed using the “Fill Holes” function, to generate a mask representing whole NeuN positive nuclei. Next, isolated DAPI stained nuclei channels were processed with corresponding NeuN masks using the image calculator function “multiply” to produce a binary image with only NeuN/DAPI double positive nuclei remaining. This image was processed in Cellpose, as mentioned above, to produce the number of NeuN/DAPI double positive cells for normalization.

### Sholl analysis

iPSC-derived neurons were cultured in geltrex coated 12 mm No.1 coverslips. Neurons were transfected with pmaxGFP^TM^ (Lonza; Morris Township, NJ) using Lipofectamine 2000 to sparsely label neurons. 24-hours post-transfection, cells were fixed with 4% PFA in PBS. Cells were stained with mouse anti-GFP (1:1000, Invitrogen, G10362) overnight at 4°C. The following day, cells were labeled with donkey anti-rabbit IgG (H+L) cross-adsorbed secondary antibody DyLight™ 488 (1:250, Invitrogen, SA5-10038). Coverslips were mounted on slides using Dako fluorescent mounting media.

Neurite morphology analyses on transfected iPSC-derived neurons were performed on randomly sampled neurons. Images of neurons were acquired using the Zeiss-880 confocal at 63X magnification with 1 Airy unit pinhole, Z-stack increments of 4.28 µm, and laser gain between 700-800 for all channels. Imaging parameters were kept consistent between inductions. Images were processed using NIH image software (ImageJ) as a maximum projection of 2x2 tiled images. Neurites were manually traced using NIH ImageJ with the SNT plugin and Sholl analysis was performed using Image J with the Sholl analysis plugin. To ensure accuracy, neurons with overlapping neurites were excluded from analysis.

### Electrophysiology

Whole-cell patch clamp recording was performed on iPSC-derived neurons 42-44 days after neural induction. Glass pipettes were filled with a solution containing the following (in mM): 135 KmeSO_3_, 5 KCl, 0.1 EGTA, 10 HEPES, 2 NaCl, 5 MgATP, 0.4 Na_2_GTP, 10 Na_2_PhosphoCreatine (pH 7.3). Cells were perfused with artificial cerebrospinal fluid (ACSF) containing (in mM)- 125 NaCl, 22.5 D-glucose, 25 NaHCO_3_, 2.5 KCl, 1.25 NaH_2_PO_4_, 3 C_3_H_3_NaO_3_, 1 C_6_H_8_O_6_, 2 CaCl, 1 MgCl, saturated with 95% O_2_, 5% CO_2_ (pH 7.3). The ACSF was continuously flowing at a rate of 2-3 ml/min at room temperature. Recording electrodes were pulled from borosilicate glass pipettes (Sutter Instrument; Novato, CA) and had a tip resistance of 3 – 5 MΩ when filled with internal solution. All recordings were performed using a MultiClamp 700B amplifier interfaced with a PC through a Digidata 1550 (Molecular Devices; Sunnyvale, CA). Data was acquired with pClamp 10.7 software (Molecular Devices; Sunnyvale, CA) and analyzed with Clampfit 10.7 (Molecular Devices; Sunnyvale, CA) and Prism (Graphpad; LaJolla; CA).

### eVLP delivery of ABE

eVLP delivery of ABE was performed by adding concentrated eVLP to HEK293T, iPSC, or iNs. For HEK293T cells, cells were dissociated into a single cell suspension with 0.25% trypsin/EDTA. 40,000 cells were plated per well of a 48-well plates. 25 µl of 100X concentrated eVLPs were added as a reverse transduction and media was changed after 24 hours. For iPSCs, cells were gently dissociated into a single cell suspension with Accutase treatment and 40,000 cells were plated on geltrex coated 48-well plates. The indicated volumes of eVLPs were treated as a reverse transduction and directly added into culturing media in either unconcentrated or concentrated form. Culturing media was changed after 24 hours. All cells were collected for analysis without selection at 72 hours. For iNs, cells were plated as normal onto glia coated cultureware. The indicated volumes of concentrated eVLPs were added as part of a standard media change with the addition of polybrene (Sigma Aldrich; Saint Louis, MO). Media was changed the following day with complete iN media with neurotrophic factors. Cells were collected for analysis without selection after 72 hours.

## Statistics

All statistical analyses were computed using Prism as described. Graphs are presented as means ± standard error of the mean (SEM), or box and whisker plots showing 10-90^th^ percentile range, with sample number indicated per experiment. Unpaired Student’s t tests, one-way ANOVA with multiple comparisons tests, and two-way ANOVA with multiple comparisons tests with Bonferroni corrections were performed to determine statistical significance at 95% confidence interval.

## Data availability statement

For original data, please contact corresponding authors, and.

## Acknowledgements

We would like to acknowledge members of the Long and Basu labs for comments and technical assistance. We would like to thank members of the NYU Langone core facilities including Cytometry and Cell Sorting Laboratory, Genome Technology Center, High Performance Computing Core, Small Instrument Fleet, and Microscopy Laboratory for their technical assistance. This work was supported by grants from the Departmental Start-Up Grant, NYU Langone Health, Kids Connect Charitable Fund, NIH (T32GM136542), NYU Clinical and Translational Science Institute (CTSI) Pilot Project (UL1TR001445, NIH/NCATS (JB/CL/OD/TB)), and Cancer Center Support Grant (P30CA016086). American Epilepsy Society Junior Investigator Award (JB), Parekh Center for Interdisciplinary Neurology (PCIN) Pilot Research Program 2023 (JB and OD), FACES (Finding a Cure for Epilepsy and Seizures), NYU Langone Health (CL/JB/OD/TB), NIH NINDS BRAIN INITIATIVE 1R01NS109994 (JB), NIH NINDS 1R01NS109362-01 (JB).

## Author contributions

C.T.K. and C.L. conceptualized the project. C.T.K., T.B., O.D., J.B., and C.L. acquired funding.

C.T.K. and N.M. performed experiments and analyzed results. T.B. and Q.Y. offered training, guidance, and oversaw experiments. V.Z., D.K., M.P. provided technical support. C.T.K. and

C.L. wrote the manuscript. C.T.K., N.M., T.B., O.D., J.B., and C.L. reviewed and edited the manuscript.

