## Supplemental figures for "Modeling and Correction of Protein Conformational Disease in iPSC-derived Neurons through Personalized Base Editing"

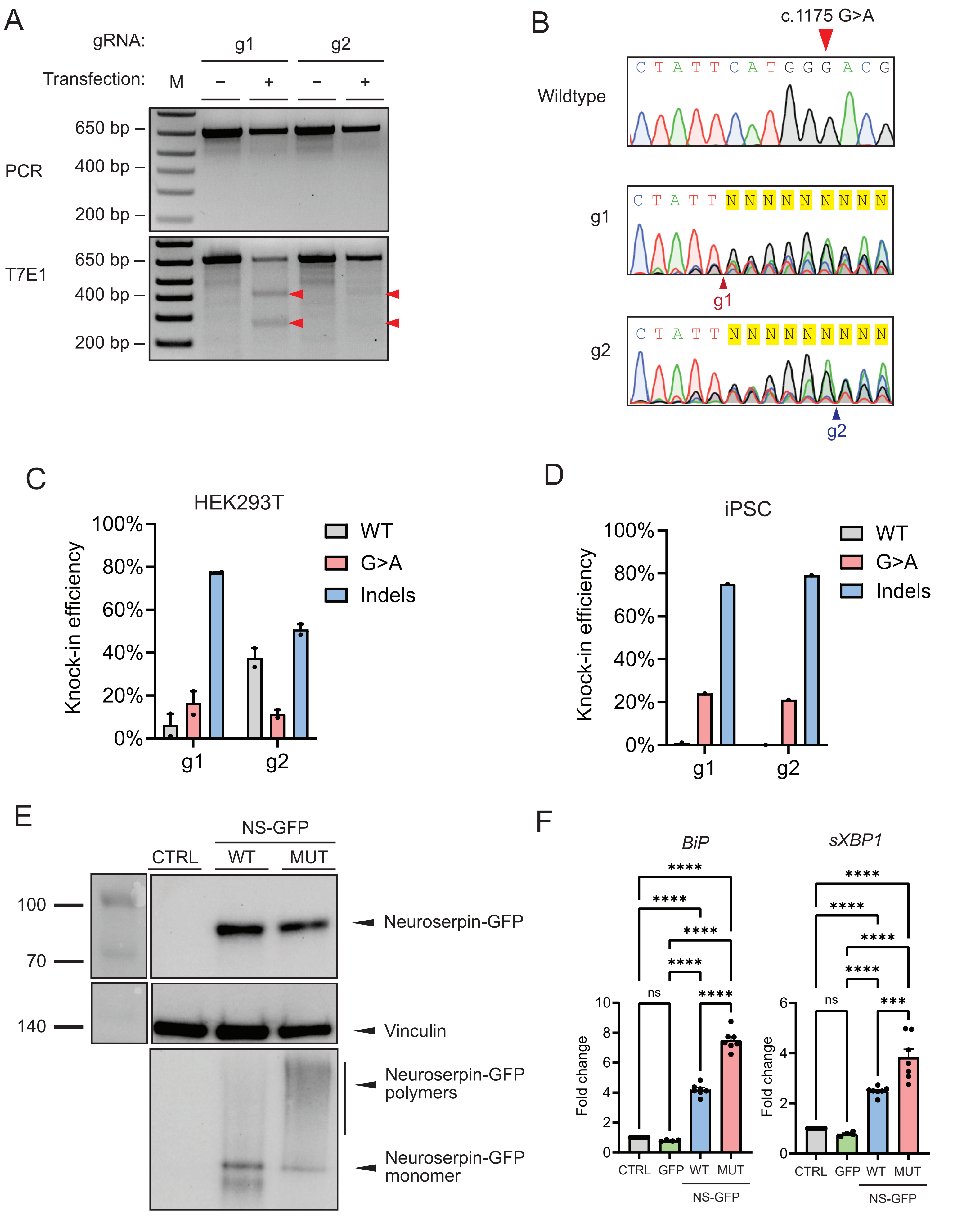

A
gRNA:
g1
g2
Transfection:
M
–
+
–
+
PCR
650 bp –
400 bp –
200 bp –
T7E1
650 bp –
400 bp –
200 bp –
B
c.1175 G>A
Wildtype
C T A T T C A T G G G A C G
g1
C T A T T N N N N N N N N N
g1
g2
C T A T T N N N N N N N N N
g2
C
HEK293T
Knock-in efficiency
100%
80%
60%
40%
20%
0%
g1
g2
WT
G>A
Indels
D
iPSC
Knock-in efficiency
100%
80%
60%
40%
20%
0%
g1
g2
WT
G>A
Indels
E
NS-GFP
CTRL
WT
MUT
100
70
140
Neuroserpin-GFP
Vinculin
Neuroserpin-GFP polymers
Neuroserpin-GFP monomer
F
BiP
sXBP1
Fold change
****
ns
***
10
8
6
4
2
0
CTRL
GFP
WT
MUT
NS-GFP


Figure S1

**Figure S1 Generation of FENIB models.**

**(A)** T7 endonuclease 1 (T7E1) cleavage assay following transfection of HEK293T cells with plasmid encoding SpCas9 and g1 or NG-SpCas9 and g2. Full intact *SERPINI1* amplicon is 650 bp. Cleavage by T7E1 results in 400 bp and 250 bp products. Red arrows indicate cleavage products. **(B)** Sanger sequencing traces of g1 and g2 treated HEK293T cells at the *SERPINI1* c.1175 G>A locus following transfection. Red and blue arrowheads indicate cleavage location of g1 and g2, respectively. **(C)** Quantification of knock-in efficiency in HEK293T cells following transfection with g1 or g2 and ssODN with *SERPINI1* c.1175 G>A variant. The graph depicts mean ± SEM and n = 2 replicates. **(D)** Quantification of knock-in efficiency in iPSC cells following transfection with g1 or g2 and ssODN with *SERPINI1* c.1175 G>A variant. The graph depicts the result of one experiment **(E)** Immunoblot of cell lysate from HEK293T cells expressing WT NS-GFP, MUT NS-GFP, or control (ctrl). Samples were resolved using anti-GFP antibody in SDS-PAGE (top), anti-vinculin antibody in SDS-PAGE as control (middle), or anti-GFP antibody in native-PAGE (bottom) to detect polymers. **(F)** Quantification of qPCR of day 7 HEK293T cells overexpressing GFP (n = 4), WT NS-GFP (n = 7), MUT NS-GFP (n = 7), or no overexpression control (n = 7) against *BiP* and spliced *XBP1* genes. (****$p<0.0001$, ***$p<0.001$, **$p<0.01$, *$p<0.5$).

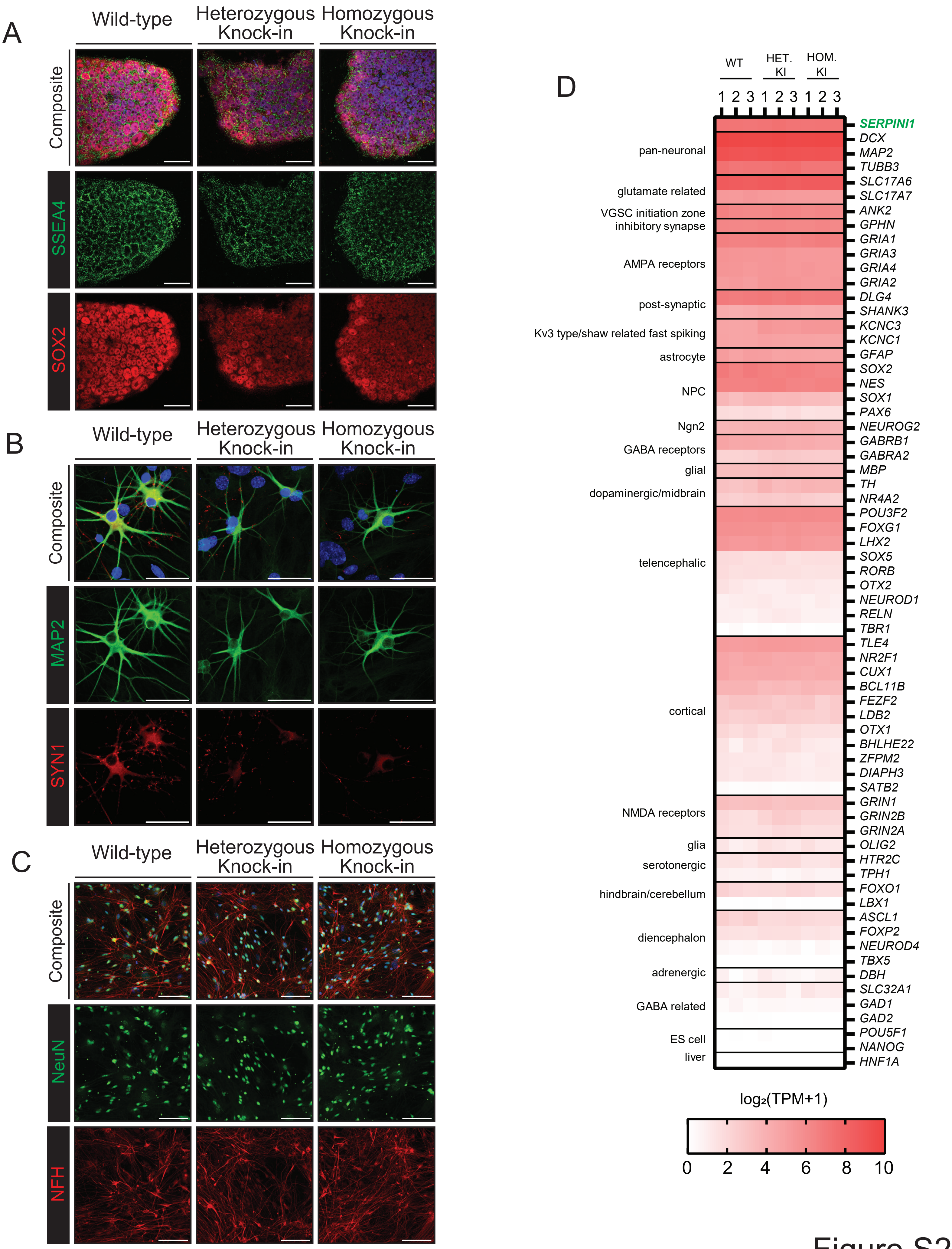
A
Wild-type
Heterozygous Knock-in
Homozygous Knock-in
Composite
SSEA4
SOX2
B
Wild-type
Heterozygous Knock-in
Homozygous Knock-in
Composite
MAP2
SYN1
C
Wild-type
Heterozygous Knock-in
Homozygous Knock-in
Composite
NeuN
NFH
D
WT
HET. KI
HOM. KI
1 2 3 1 2 3 1 2 3
SERPINI1
pan-neuronal
DCX
MAP2
TUBB3
glutamate related
SLC17A6
SLC17A7
VGSC initiation zone
ANK2
inhibitory synapse
GPHN
AMPA receptors
GRIA1
GRIA3
GRIA4
GRIA2
post-synaptic
DLG4
SHANK3
Kv3 type/shaw related fast spiking
KCNC3
KCNC1
astrocyte
GFAP
NPC
SOX2
NES
SOX1
PAX6
Ngn2
NEUROG2
GABA receptors
GABRB1
GABRA2
glial
MBP
dopaminergic/midbrain
TH
NR4A2
telencephalic
POU3F2
FOXG1
LHX2
SOX5
RORB
OTX2
NEUROD1
RELN
TBR1
cortical
TLE4
NR2F1
CUX1
BCL11B
FEZF2
LDB2
OTX1
BHLHE22
ZFPM2
DIAPH3
SATB2
NMDA receptors
GRIN1
GRIN2B
GRIN2A
glia
OLIG2
serotonergic
HTR2C
TPH1
hindbrain/cerebellum
FOXO1
LBX1
diencephalon
ASCL1
FOXP2
NEUROD4
TBX5
adrenergic
DBH
GABA related
SLC32A1
GAD1
GAD2
ES cell
POU5F1
NANOG
liver
HNF1A
log₂(TPM+1)
0 2 4 6 8 10

Figure S2

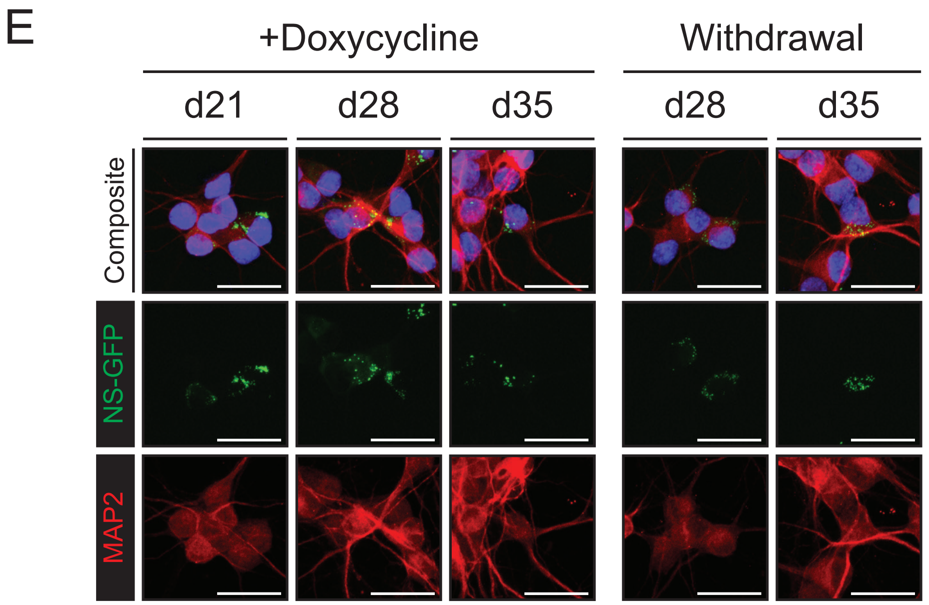
E
+Doxycycline
Withdrawal
d21
d28
d35
d28
d35
Composite
NS-GFP
MAP2

Figure S2

**Figure S2 Characterization of iPSC-derived neurons.**

**(A)** Representative confocal microscopy images of iPSCs labeled with antibodies specific for stage-specific embryonic antigen 4 (SSEA4) (green) and SRY-Box transcription factor 2 (SOX2) (red). Scale bar, 50 µm. **(B)** Representative confocal microscopy images of d21 neurons labeled with antibodies for neuron specific microtubule associated protein 2 (MAP2) (green) and synapse protein synapsin I (SYN1) (red). Scale bar, 50 µm. **(C)** Representative confocal microscopy of day 35 neurons. Cells stained with antibodies for neuron cytoskeleton protein neurofilament heavy chain (NFH) (green) and neuronal nuclei protein (NeuN) (red). Scale bar, 100 µm. **(D)** Heatmap of *SERPINI1* (green) and neuronal genes selected from Zhang et al. (2013). Expression level was calculated by $\log_2$ of transcripts per million (TPM) + 1 and is depicted by color as indicated on bottom. **(E)** Representative confocal microscopy images of MUT NS-GFP neurons labeled with MAP2 (red) and DAPI (blue), harboring NS-GFP aggregates (green). Scale bar, 25 µm.

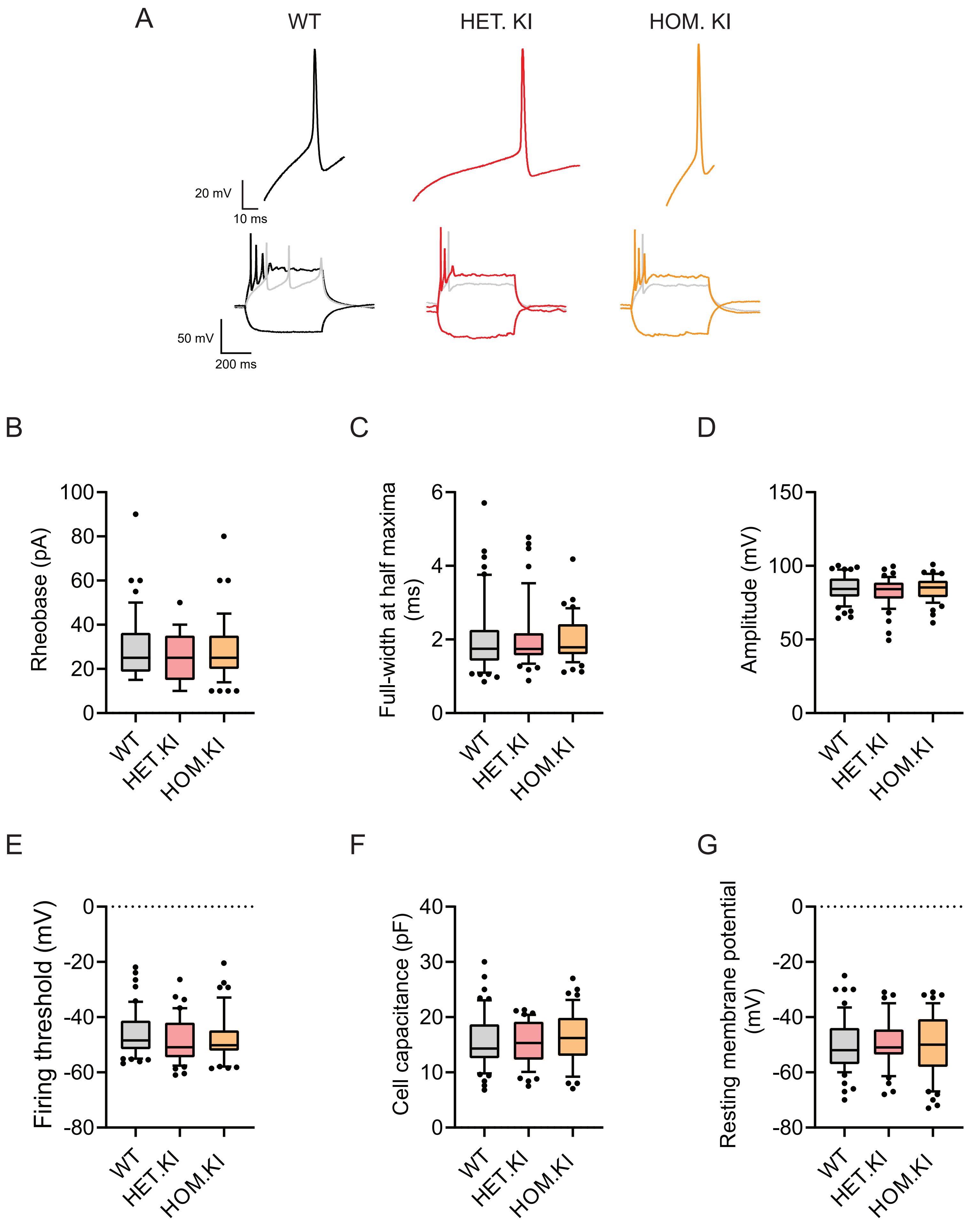
A
WT
HET. KI
HOM. KI
20 mV
10 ms
50 mV
200 ms
B
C
D
Rheobase (pA)
Full-width at half maxima (ms)
Amplitude (mV)
E
F
G
Firing threshold (mV)
Cell capacitance (pF)
Resting membrane potential (mV)
WT
HET.KI
HOM.KI

Figure S3

**Figure S3 Evoked action potential analysis in KI neurons.**

**(A-F)** Whole cell current patch clamp of day 42-44 KI FENIB neurons to assess evoked APs. The graphs depict individual data points and box and whisker plot displaying 10-90 percentile range and n = 46-54 neurons (WT = 52, HET. KI = 46 HOM. KI = 47, WT = 54 only for rheobase). **(A)** Quantification of rheobase. **(B)** Quantification of full-width at half maxima. **(C)** Quantification of amplitude. **(D)** Quantification of firing threshold. **(E)** Quantification of cell capacitance. **(G)** Quantification of resting membrane potential. WT= wild-type neurons, HET. KI = heterozygous knock-in neurons, HOM. KI = homozygous knock-in neurons.

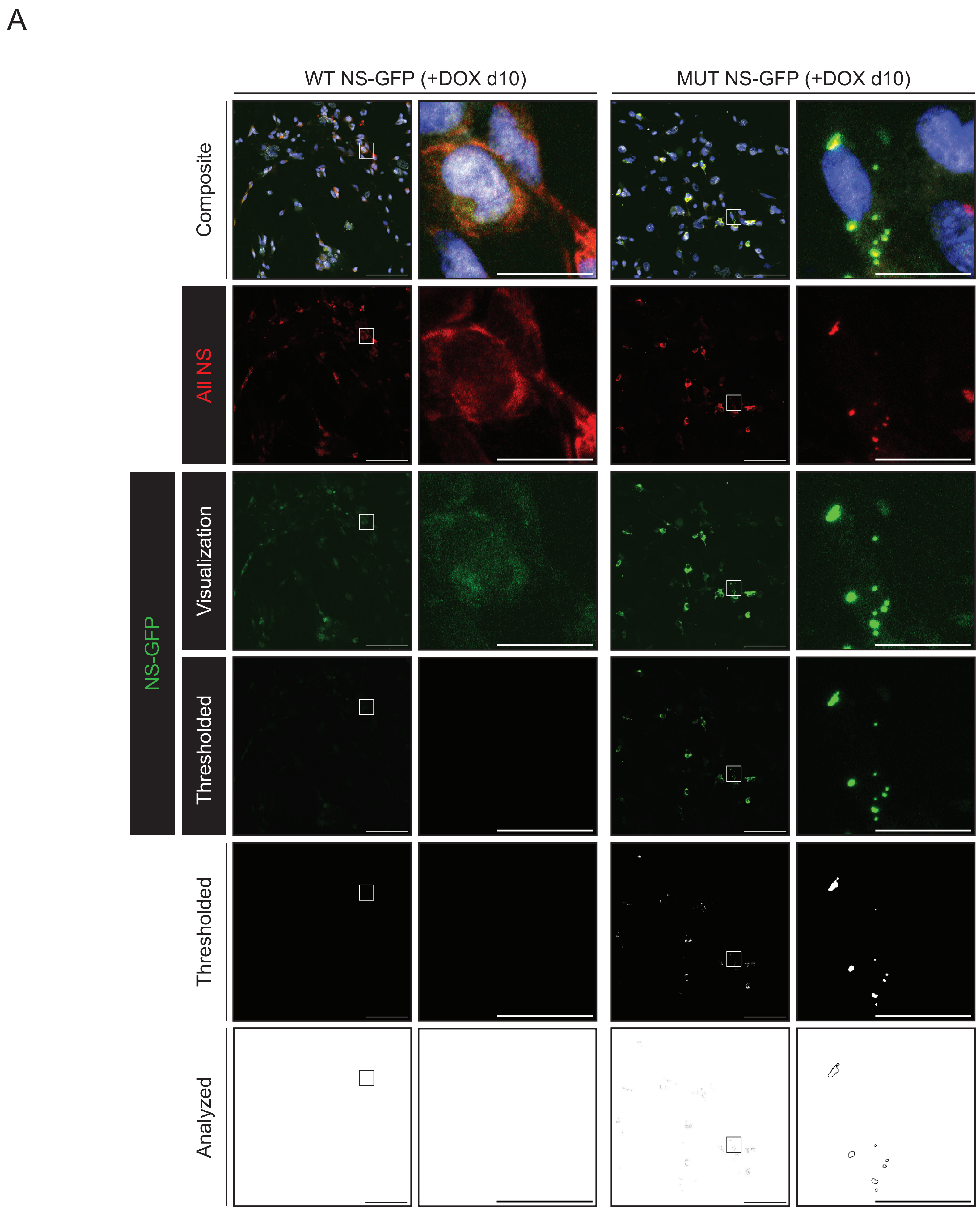

A
WT NS-GFP (+DOX d10)
MUT NS-GFP (+DOX d10)
Composite
All NS
NS-GFP
Visualization
Thresholded
Thresholded
Analyzed


Figure S4

**Figure S4 Overview of NS-GFP aggregate quantification.**

**(A)** Representative overview of NS-GFP aggregate analysis of MUT NS-GFP +DOX d10 and WT NS-GFP +DOX d10 iNs. Cells expressing WT or MUT NS-GFP (green) are labeled with antibodies specific for all neuroserpin forms (all NS) (red) and NeuN (white) and are stained with DAPI (blue). Each sample is shown as a maximal projected, 2x2 titled image at 63X magnification (left) or as a focused inset (white box, right). Scale bar = 100 μm, inset scale bar = 20 μm. Images labeled, "NS-GFP Visualization" have fluorescence contrast evenly increased to aid visualization. Images labeled, "NS-GFP Thresholded" have fluorescence contrast set at the fluorescence threshold. Images labeled "Thresholded" have NS-GFP signal assigned as either "diffuse" or "aggregate" based on pixel intensity. All pixels with fluorescent signal below the threshold are determined to be diffuse signal and are assigned a value of 0 (black). All pixels with fluorescence signal above the threshold are determined to be aggregates and are assigned a value of 255 (white). Images labeled "Analyzed" measure the number of aggregates and area of aggregates as assigned by the threshold.

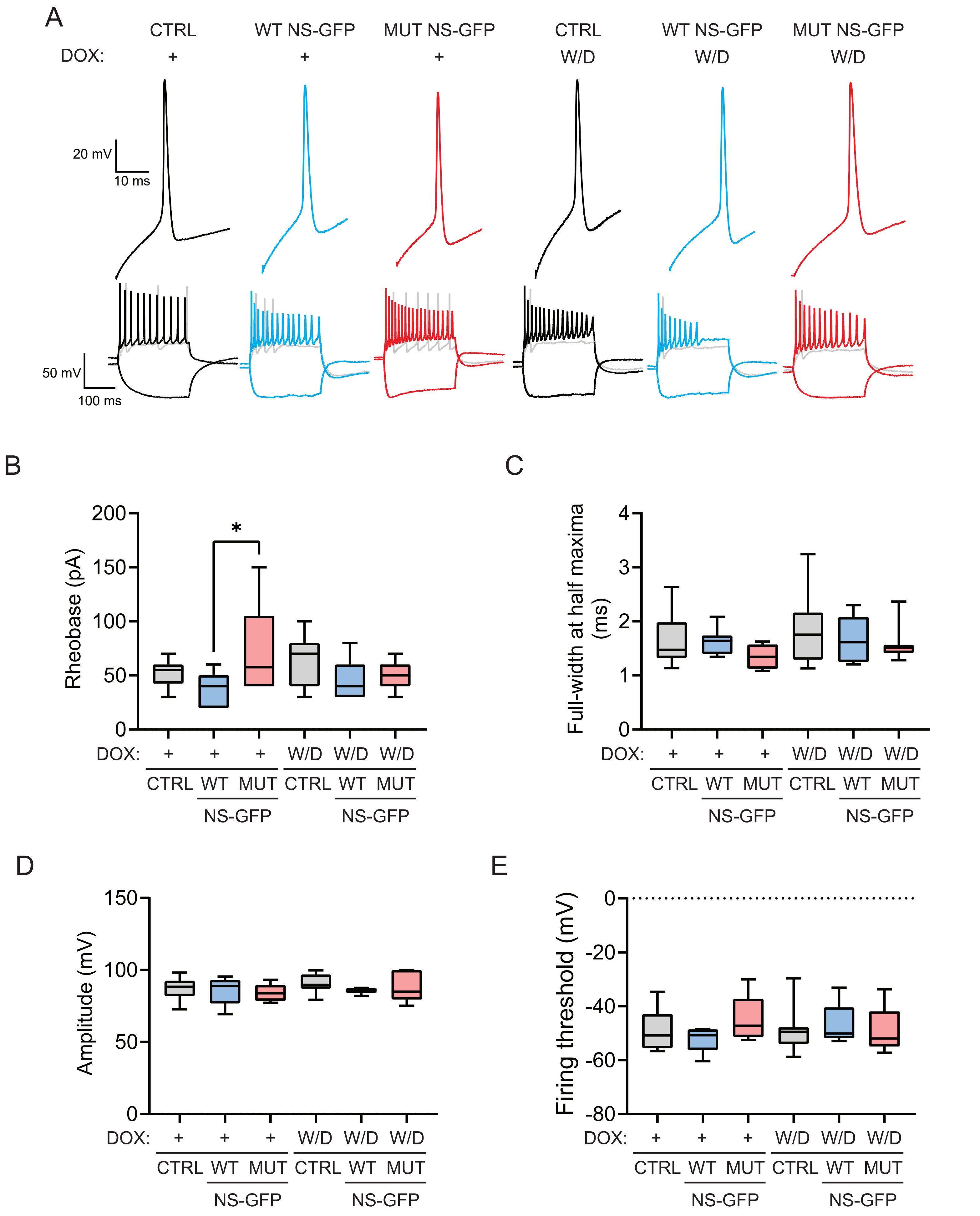
A
CTRL
WT NS-GFP
MUT NS-GFP
CTRL
WT NS-GFP
MUT NS-GFP
DOX: + + + W/D W/D W/D
20 mV
10 ms
50 mV
100 ms
B
Rheobase (pA)
200
150
100
50
0
*
DOX: + + + W/D W/D W/D
CTRL WT MUT CTRL WT MUT
NS-GFP NS-GFP
C
Full-width at half maxima (ms)
4
3
2
1
0
DOX: + + + W/D W/D W/D
CTRL WT MUT CTRL WT MUT
NS-GFP NS-GFP
D
Amplitude (mV)
150
100
50
0
DOX: + + + W/D W/D W/D
CTRL WT MUT CTRL WT MUT
NS-GFP NS-GFP
E
Firing threshold (mV)
0
-20
-40
-60
-80
DOX: + + + W/D W/D W/D
CTRL WT MUT CTRL WT MUT
NS-GFP NS-GFP

Figure S5

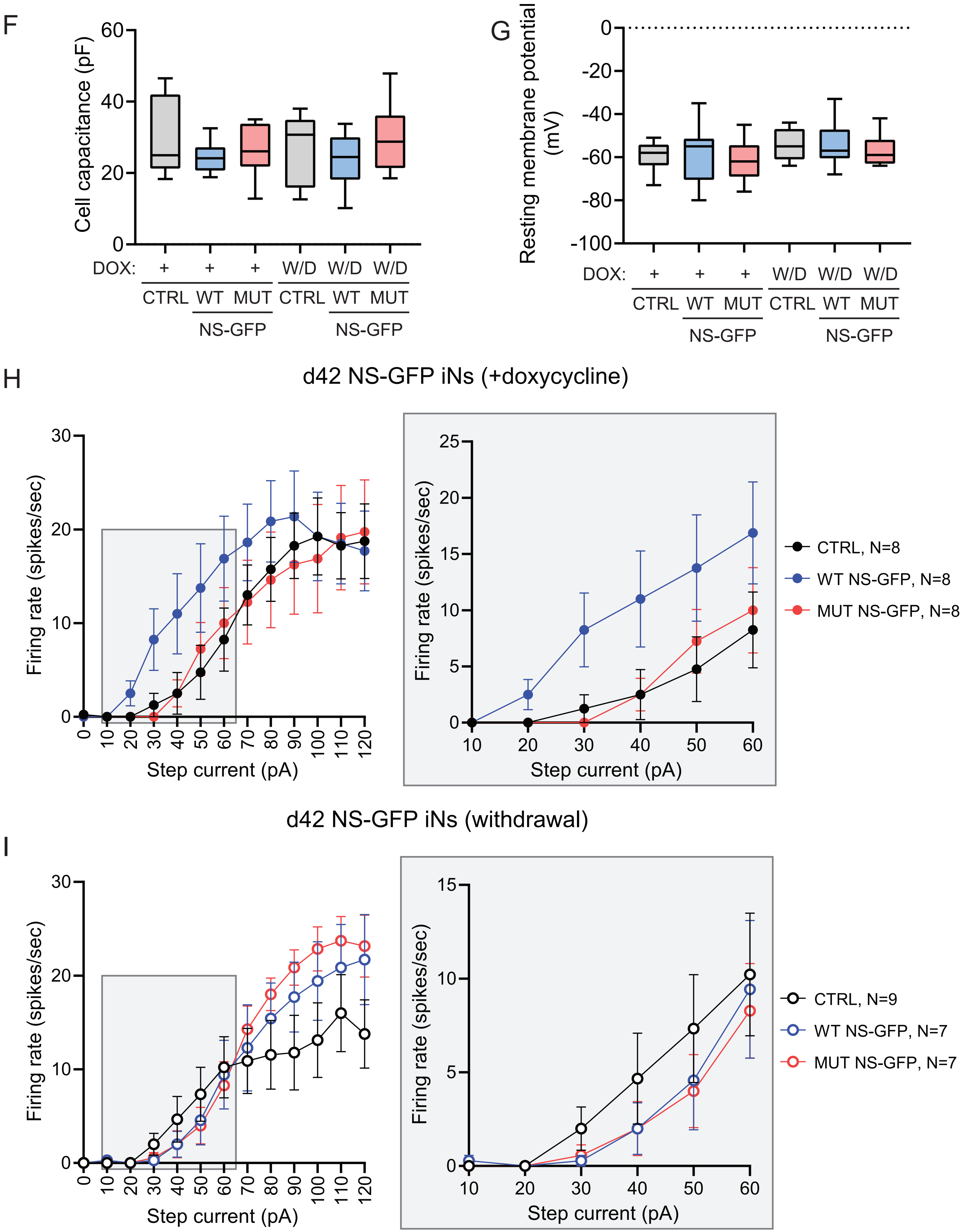
F
Cell capacitance (pF)
60
40
20
0
DOX: + + + W/D W/D W/D
CTRL WT MUT CTRL WT MUT
NS-GFP NS-GFP
G
Resting membrane potential (mV)
0
-20
-40
-60
-80
-100
DOX: + + + W/D W/D W/D
CTRL WT MUT CTRL WT MUT
NS-GFP NS-GFP
H
d42 NS-GFP iNs (+doxycycline)
Firing rate (spikes/sec)
30
20
10
0
0 10 20 30 40 50 60 70 80 90 100 110 120
Step current (pA)
25
20
15
10
5
0
10 20 30 40 50 60
Step current (pA)
CTRL, N=8
WT NS-GFP, N=8
MUT NS-GFP, N=8
d42 NS-GFP iNs (withdrawal)
I
Firing rate (spikes/sec)
30
20
10
0
0 10 20 30 40 50 60 70 80 90 100 110 120
Step current (pA)
15
10
5
0
10 20 30 40 50 60
Step current (pA)
CTRL, N=9
WT NS-GFP, N=7
MUT NS-GFP, N=7

Figure S5

**Figure S5 Evoked action potential analysis in NS-GFP neurons.**

**(A-G)** Whole cell current patch clamp of day 42-44 NS-GFP neurons with continuous doxycycline or doxycycline withdrawal at day 21 to assess evoked APs. **(A)** Representative single action potentials and action potential trains. Single action potentials are the first evoked action potential taken at rheobase. Representative action potential trains are shown at rheobase in grey and two times rheobase in color (black for ctrl, blue for WT NS-GFP, and red for MUT NS-GFP). **(B-G)** The graphs depict box and whisker plot displaying 10-90 percentile range and n = 7-9 neurons (CTRL +DOX = 8, WT NS-GFP +DOX = 8, MUT NS-GFP +DOX = 8, CTRL W/D = 9, WT NS-GFP W/D = 7, MUT NS-GFP W/D = 7). **(B)** Quantification of rheobase. **(C)** Quantification of full-width at half maxima. **(D)** Quantification of amplitude. **(E)** Quantification of firing threshold. **(F)** Quantification of cell capacitance. **(G)** Quantification of resting membrane potential. **(H)** Quantification of firing rate of d42 NS-GFP neurons with continuous NS-GFP expression (+doxycycline) with an inset focusing on the 10-60 pA region. Inset is indicated via a grey box. **(I)** Quantification of firing rate of d42 NS-GFP neurons with NS-GFP expression withdrawn at d21 (withdrawal) with inset focusing on the 10-60 pA region. Inset is indicated via a grey box.

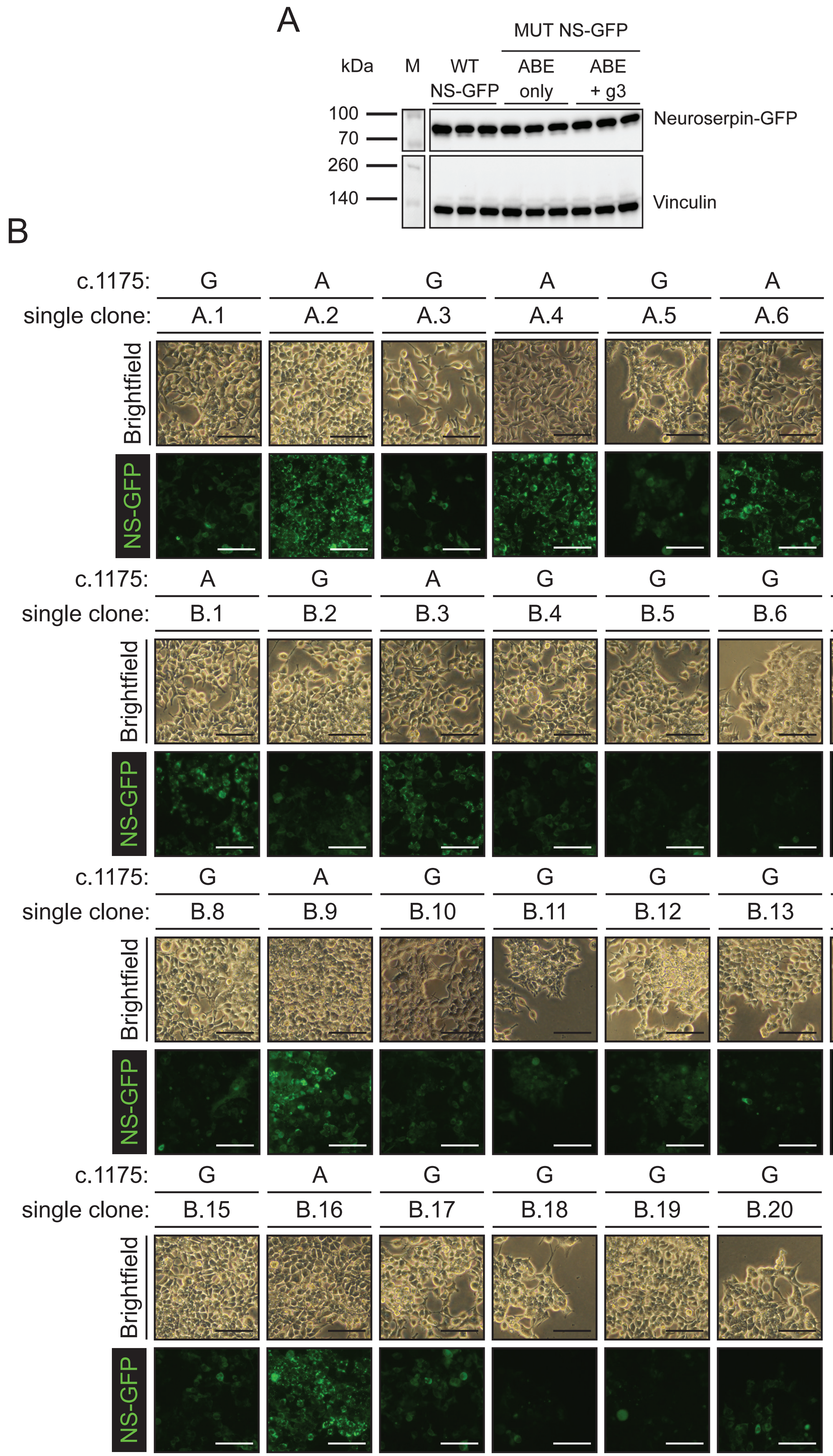
A
MUT NS-GFP
kDa
M
WT
NS-GFP
ABE
only
ABE
+ g3
100
70
260
140
Neuroserpin-GFP
Vinculin
B
c.1175: G A G A G A
single clone: A.1 A.2 A.3 A.4 A.5 A.6
Brightfield
NS-GFP
c.1175: A G A G G G G
single clone: B.1 B.2 B.3 B.4 B.5 B.6 B.7
Brightfield
NS-GFP
c.1175: G A G G G G G
single clone: B.8 B.9 B.10 B.11 B.12 B.13 B.14
Brightfield
NS-GFP
c.1175: G A G G G G
single clone: B.15 B.16 B.17 B.18 B.19 B.20
Brightfield
NS-GFP

Figure S6

**Figure S6 Correction of MUT NS-GFP in HEK293Ts allows for aggregate clearance.**

**(A)** Immunoblot of cell lysate from WT NS-GFP, ABE only MUT NS-GFP, or ABE + g3 treated MUT NS-GFP HEK293T cells. Samples were resolved using anti-neuroserpin antibody (top), and anti-vinculin antibody as a loading control (bottom). **(B)** Representative fluorescent and brightfield microscopy of clonally expanded ABE + g3 treated MUT NS-GFP HEK293T cells. Clones derived from MUT NS-GFP clones as follows: Clone A (A.1-A.6) and from Clone B (B.1-B20). Genotype at variant c.1175 G>A is written above each clone name (G = WT, A = MUT). Scale bar, 100 μm.

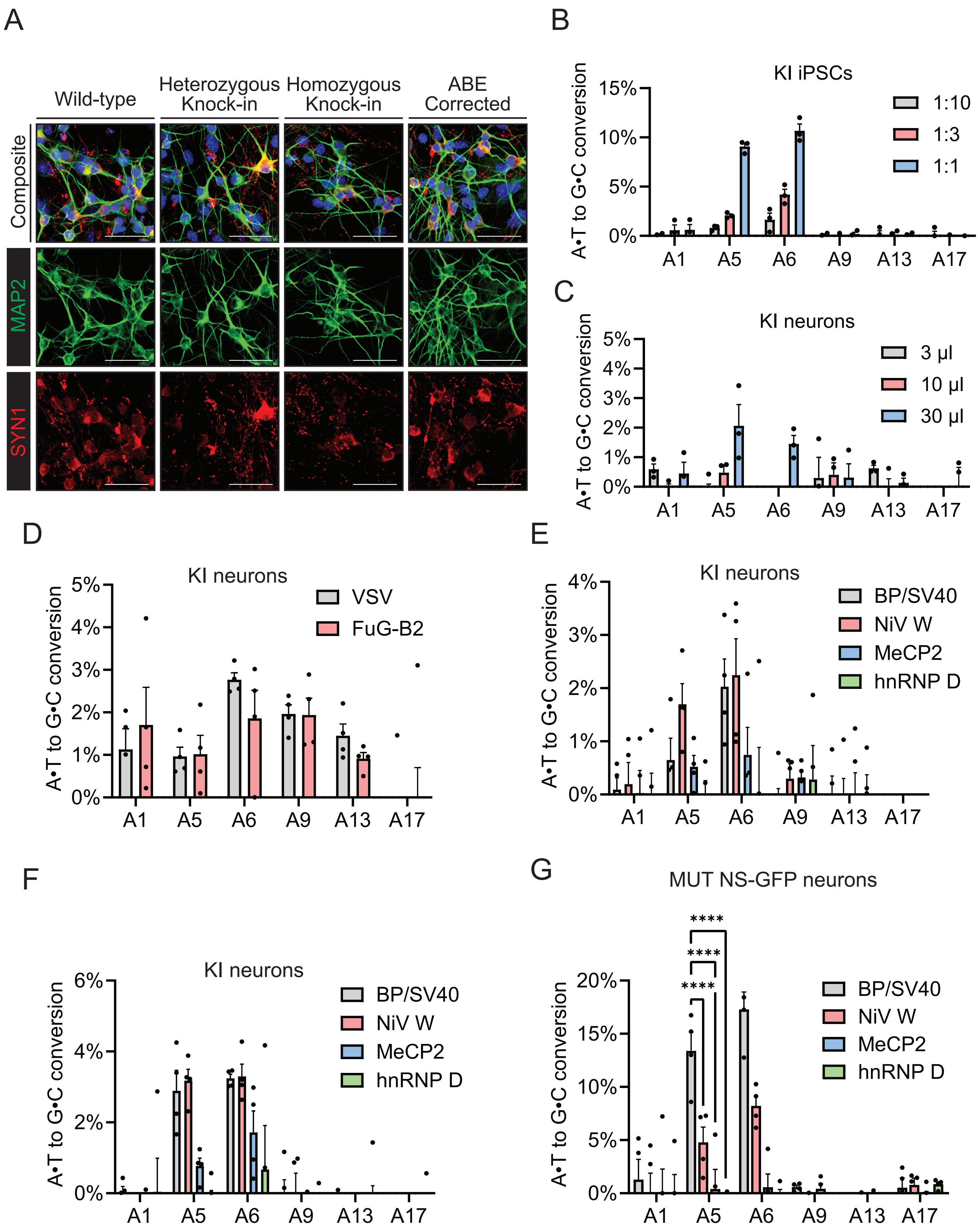
A
Wild-type
Heterozygous Knock-in
Homozygous Knock-in
ABE Corrected
Composite
MAP2
SYN1
B
KI iPSCs
1:10
1:3
1:1
A•T to G•C conversion
C
KI neurons
3 μl
10 μl
30 μl
D
KI neurons
VSV
FuG-B2
E
KI neurons
BP/SV40
NiV W
MeCP2
hnRNP D
F
KI neurons
G
MUT NS-GFP neurons
****
A1
A5
A6
A9
A13
A17

Figure S7

**Figure S7 eVLP editing in iPSC and iPSC-derived neurons.**

**(A)** Representative confocal microscopy images of d35 neurons labeled with antibodies specific for MAP2 (green) and SYN1 (red). Scale bar, 50 µm. **(B-G)** Quantification of eVLP editing. The graphs depict mean ± SEM of n = 3 replicates. **(B)** Quantification of eVLP-ABE editing of *SERPINI1* c1175 G>A variant in HOM. KI iPSCs using unconcentrated eVLP supernatant in ratios of 1:10, 1:3, and 1:1 (unconcentrated eVLP supernatant: Stemflex media). **(C)** Quantification of concentrated eVLP editing in day 7 HOM. KI neurons. **(D)** Quantification of concentrated eVLP editing in day 7 HOM. KI neurons testing VSV-G and FuG-B2 glycoproteins. **(E)** Quantification of concentrated eVLP editing in day 7 HOM. KI neurons testing classical NLS BP/SV40, or nonclassical NLS NiV W, MeCP2, and hnRNPD tagged MMLV-gag fused ABEs with VSV-G glycoprotein eVLPs. **(F)** Quantification of concentrated eVLP editing in day 7 HOM. KI neurons testing classical NLS BP/SV40, or nonclassical NLS NiV W, MeCP2, and hnRNPD tagged MMLV-gag fused ABEs with FuG-B2 glycoprotein eVLPs. **(G)** Quantification of concentrated eVLP editing in day 7 MUT NS-GFP neurons testing classical NLS BP/SV40, or nonclassical NLS NiV W, MeCP2, and hnRNPD tagged MMLV-gag fused ABEs with VSV-G glycoprotein eVLPs in the presence of 5 µg/ml polybrene.
